# AL-PHA beads: bioplastic-based protease biosensors for global health applications

**DOI:** 10.1101/2020.06.18.159921

**Authors:** Richard J. R. Kelwick, Alexander J. Webb, Yizhou Wang, Amelie Heliot, Fiona Allan, Aidan M. Emery, Michael R. Templeton, Paul S. Freemont

## Abstract

Proteases are multi-functional proteolytic enzymes that have complex roles in human health and disease. Therefore, the development of protease biosensors can be beneficial to global health applications. To this end, we developed Advanced proteoLytic detector PolyHydroxyAlkanoates (AL-PHA) beads – a library of over 20 low-cost, biodegradable, bioplastic-based protease biosensors. Broadly, these biosensors utilise PhaC-reporter fusion proteins that are bound to microbially manufactured polyhydroxyalkanoate beads. In the presence of a specific protease, superfolder green fluorescent reporter proteins are cleaved from the AL-PHA beads - resulting in a loss of bead fluorescence. The Tobacco Etch Virus (TEV) AL-PHA biosensor detected the proteolytic activity of at least 1.85 pM of AcTEV. AL-PHA beads were also engineered to detect cercarial elastase from *Schistosoma mansoni*-derived cercarial transformation fluid (SmCTF) samples, as well as cancer-associated metalloproteinases in extracellular vesicle and cell-conditioned media samples. We envision that AL-PHA beads could be further developed for use in resource-limited settings.

## Introduction

Synthetic biology is an established scientific field based upon engineering design principles, that has led to innovations in the development of biosensors and bioreporters geared towards global health applications [1,2]. These diverse applications have inspired a multitude of biosensor designs that have innovated beyond typical electrochemical formats, towards whole-cell bioreporters (WCBs) and cell-free biosensors [3]. More recently, convergences between the materials sciences and synthetic biology are opening up new opportunities for global health biosensor applications [4]. In particular, we envisage that functionalised biomaterials may enable the emergence of novel strategies for detecting biomedically important proteases [5]. Proteases are multi-functional proteolytic enzymes, that have complex roles in human health and disease [6]. Protease functions are diverse and can be broad, such as aiding food digestion, or highly evolved and specialised targeting more specific substrates [6]. Exemplars of proteases that have evolved to serve complex biological functions can be found within the matrix metalloproteinase (MMP), the A Disintegrin And Metalloproteinase (ADAM) and A Disintegrin and Metalloproteinase with Thrombospondin motifs (ADAMTS) protease families [7]. Members of these protease families contribute to an array of biological processes including: cellular metabolism, cell-signalling, cell-migration, immunomodulation and tissue remodelling [8,9]. Changes in human metalloproteinase gene expression and/or their proteolytic activities can lead to cardiovascular or inflammatory pathologies, neurodegenerative diseases, changes in immunoregulation and cancer [10,11]. Proteases also have important roles in communicable diseases, whereby infectious microorganisms and parasites employ proteases to support pathogenesis [12]. In the case of schistosomiasis (also known as bilharzia or snail fever), a neglected tropical disease that affects over 250 million people worldwide [13–15], the invasive *Schistosoma* cercariae release a cocktail of proteases, including elastase, that help the parasite invade their host through the skin [16,17].

Understanding the activities of proteases can lead to important insights into communicable and non-communicable diseases [6]. Indeed, novel protease detection strategies, especially those intended for field use, may be beneficial to many different clinical, biotechnological, environmental and epidemiological global health applications [5,18–21]. For example, in our previous study we used a synthetic biology approach to engineer *Escherichia coli* and *Bacillus subtilis* WCBs that can detect the elastase activity from the cercariae of the parasite *Schistosoma mansoni* [22]. Importantly, our study demonstrated the detection of the proteolytic activity of a specific protease (i.e. cercarial elastase) within complex biological samples. However, the implementation of WCBs within global health settings is challenging and many other complex practical, cultural, societal, data protection and regulatory concerns must also be addressed [23]. Understandably, amongst those concerns, the accidental release of living engineered WCBs is commonly cited [24]. In response to these challenges, the development of WCBs has led to important innovations in physical (e.g. sealing WCBs within devices) and genetic (e.g. genetically encoded kill switches or auxotrophy) containment strategies, that help mitigate the risk of accidental release [23]. Whilst these biological containment strategies are impressive, we anticipate that non-living biosensors may be desirable in certain contexts.

To this end, we developed modular, functionalised, polyhydroxyalkanoates (PHAs)-based, bioplastic beads for protease detection. We termed these biosensor beads - **A**dvanced proteo**L**ytic detector **PHA**s (**AL-PHA**) beads and initially optimised their design using a commercially available Tobacco Etch Virus (TEV) protease. As a proof-of-concept for a global health applications, AL-PHA biosensors were assayed against *S. mansoni* derived samples containing soluble cercarial antigens, termed cercarial transformation fluid (SmCTF). AL-PHA biosensors successfully detected cercarial elastase activity within these samples. We also showed that AL-PHA beads engineered to detect cancer-associated metalloproteinases including: MMPs, ADAMs and ADAMTSs were functional. Most notably, AL-PHA beads detected recombinant MMP14 and extracellular vesicle (EV)-associated ADAM10 derived from an *in vitro* model of non-small cell lung cancer (nsclc). Furthermore, we also demonstrate the potential use of AL-PHA beads in a high-throughput screening context - whereby an entire library of metalloproteinase biosensors was tested in parallel. To the best of our knowledge, this proof-of-concept study is the first to demonstrate the use of functionalised PHAs-based protease biosensors for health care applications in resource-limited environments.

## Results

### Functionalised polyhydroxyalkanoates (PHAs)-based beads for protease detection

PHAs represent a diverse family of biopolymers with different mechanical characteristics, thermal properties, biodegradabilities and biocompatibilities [25]. Poly-3-hydroxybutyrate (P(3HB)) is one of the most well studied PHAs polymers and can be used within food packaging or for other industrial applications, including medical and tissue engineering applications [26]. Polymerised P(3HB) naturally forms into spherical granules (beads) within suitably engineered *E. coli* [27]. Interestingly, these PHAs beads can also be functionalised *in vivo* with engineered fusion proteins enabling novel protein purification strategies [28], as well as the development of PHAs-based vaccines [29]. To further extend the applications of functionalised PHAs materials we engineered our own PhaC-fusion proteins, which could act as protease biosensors, on the surfaces of these PHAs beads – which we termed AL-PHA beads (Fig. 1).

**Figure 1.**
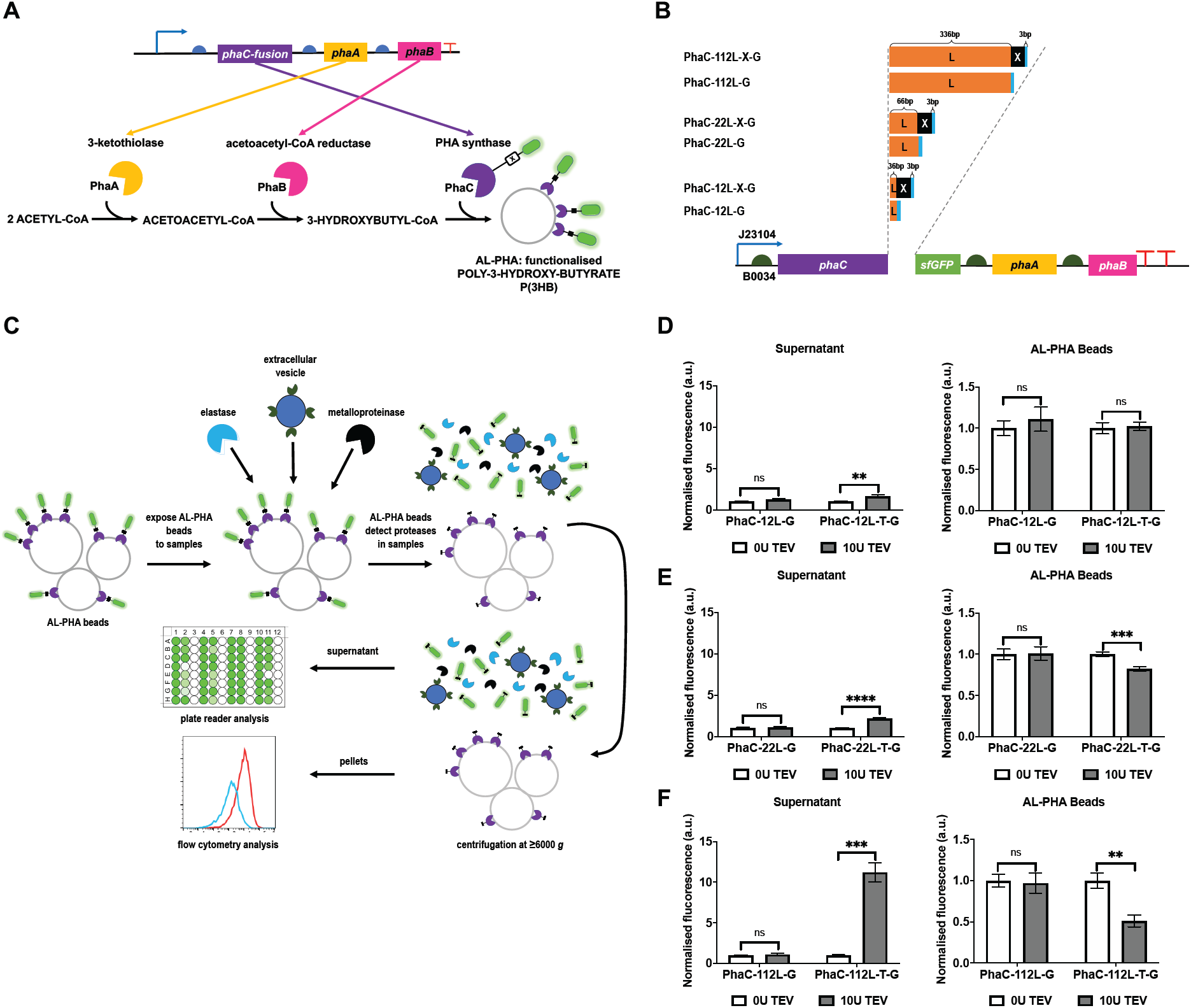
AL-PHA biosensor design optimisation. **(A)** Schematic of the *phaCAB* biosynthetic operon and the enzymatic pathway used for production of AL-PHA biosensor beads in engineered *Escherichia coli*. Initially, acetyl-CoA is processed by PhaA (3-ketothiolase) to form acetoacetyl-CoA. Subsequently, PhaB (acetoacetyl-CoA reductase) reduces acetoacetyl-CoA to form the monomer (*R*)-3-hydroxybutyl-CoA ((*R*)-3HB-CoA). Finally, (*R*)-3HB-CoA is polymerised by PhaC (PHA synthase) to form the final PHAs polymer - P(3HB). This P(3HB) polymer naturally forms into spherical granules (beads), that display PhaC-fusion proteins, within suitably engineered *E. coli*. **(B)** Schematic of engineered *phaCAB* operons for the production of control and protease detection AL-PHA biosensors. Abbreviations: 12L, 22L and 112L denote the amino acid length of the flexible linker, X denotes the protease recognition motif site for either TEV (T), cercarial elastase (E) MMP (P), ADAM (M) or ADAMTS (TS) proteases and G denotes superfolder green fluorescent protein (sfGFP). **(C)** AL-PHA biosensor assay workflow. Analysis of 12L **(D)**, 22L **(E)** and 112L **(F)** control and TEV protease AL-PHA biosensors. AL-PHA biosensors were treated with either 0 units (0 U) or 10 units (10 U) of AcTEV protease. Proteolytically released sfGFP in supernatant samples were analysed using a CLARIOstar plate reader (483-14 nm/530-30nm) and these fluorescence data were normalised against untreated controls of the same biosensor batch. AL-PHA beads were analysed using flow cytometry and AL-PHA bead geometric mean (BL1-A, 488nm/530-30nm) of AcTEV treated beads were normalised against untreated controls of the same biosensor batch. Error bars denote standard error of the mean, n=4-8, Student *t*-test **P<0.01, ***P<0.001, ****P<0.0001or not statistically significant (ns).

To achieve this, we adapted a highly active C104 *phaCAB* biosynthetic operon from our previous studies, to develop AL-PHA producing operons in engineered *E. coli* [30,31]. These AL-PHA operons contain PhaC-fusion proteins have been designed to incorporate a flexible and modular amino acid linker that comprises protease-specific cleavage sites, and a GFP (sfGFP) reporter protein (Fig. 1A & 1B; Supplementary Tables 1-4; Supplementary Fig. 1). By changing the protease cleavage sites, we were able to construct a suite of AL-PHA bead protease biosensors (Supplementary Fig. 2; Supplementary Table 1). These PhaC-fusion proteins were designed such that specific proteases can be detected via the proteolytic cleavage of sfGFP from the surface of the AL-PHA beads.

AL-PHA beads were produced in *E. coli* cells, isolated using a sonication-based method and tested using a simple assay (Fig. 1C). Essentially, control or protease specific AL-PHA beads were incubated with proteases, with optimal reaction conditions tailored to the specific protease being tested (See materials and methods). Post-incubation, AL-PHA beads were centrifuged allowing the supernatant and pellet to be analysed separately. This allows proteolytically released sfGFP in the supernatant to be measured using a plate reader, whilst the associated concomitant loss in AL-PHA bead fluorescence was measured using flow cytometry.

To assess whether our PhaC-sfGFP fusion proteins were correctly localised on the AL-PHA bead surface, we used the non-specific protease trypsin. As expected, trypsin treatment (1 μg) significantly decreased AL-PHA bead fluorescence compared to untreated controls (0 μg; Supplementary Fig. 3). Usefully, this reduction in bead fluorescence was observable to the naked eye when the beads were pelleted and placed onto a transilluminator (Supplementary Fig. 3C). This initial data supported the notion that the engineered PhaC-fusion proteins are accessible and susceptible to proteolytic activity on the surface of the AL-PHA beads.

We next chose three different proteolytic linker designs specific for TEV protease, which differed only in length, as follows: 12 amino acids (12L), twenty-two amino acids (22L) and one hundred and twelve amino acids (112L) (Fig. 1B; Supplementary Fig. 1; Supplementary Table 2). We previously identified that longer linker lengths can increase proteolytic sensitivity, likely through improving cleavage site access [22]. AL-PHA beads with the relevant TEV linker designs were assayed with either 0 or 10 U of AcTEV protease and incubated at 30°C for 2 hours with shaking. Post-assay supernatants and AL-PHA beads were assessed, separately, using either plate reader (supernatant) or flow cytometry (AL-PHA bead) workflows (Supplementary Fig. 4). In comparison to untreated controls (0 U AcTEV), supernatant fluorescence levels increased for all AcTEV treated (10 U) AL-PHA bead samples indicating that all three linker designs can specifically detect AcTEV protease (Fig. 1D-F). Interestingly, the most sensitive linker was 112L (PhaC-112L-T-G), with a ∼10-fold relative increase in supernatant fluorescence levels compared to controls (Fig. 1F), which we quantified as being equivalent to 1.87 μM sfGFP being released (Supplementary Fig. 5). By analysing flow cytometry data (Fig. 1F; Supplementary Fig. 5) we estimated that ∼187 pmoles of GFP is being released from ∼300,000 AL-PHA beads, which is within the range of previous studies in terms of surface bead coverage [28,32]. Furthermore, the concomitant reduction in AL-PHA bead fluorescence was also significantly more pronounced for 112L (∼49% decrease; Fig. 1F) and this reduction was observable to the naked eye using a transilluminator (Supplementary Fig. 6). We also observed that after 2 hours of treatment the PhaC-112L-T-G AL-PHA beads are sufficiently sensitive to detect 0.5 U of AcTEV activity (Fig. 2A, B), which is ∼1.85 pM of AcTEV protease. Taken together, our data shows that AL-PHA beads with the longest linker (112L) are the most sensitive biosensor design. Furthermore, AL-PHA beads are robust and remain visibly fluorescent for at least 12 months when stored at 4 °C (Supplementary Fig. 7), which is useful for downstream field and point-of-care applications.

**Figure 2.**
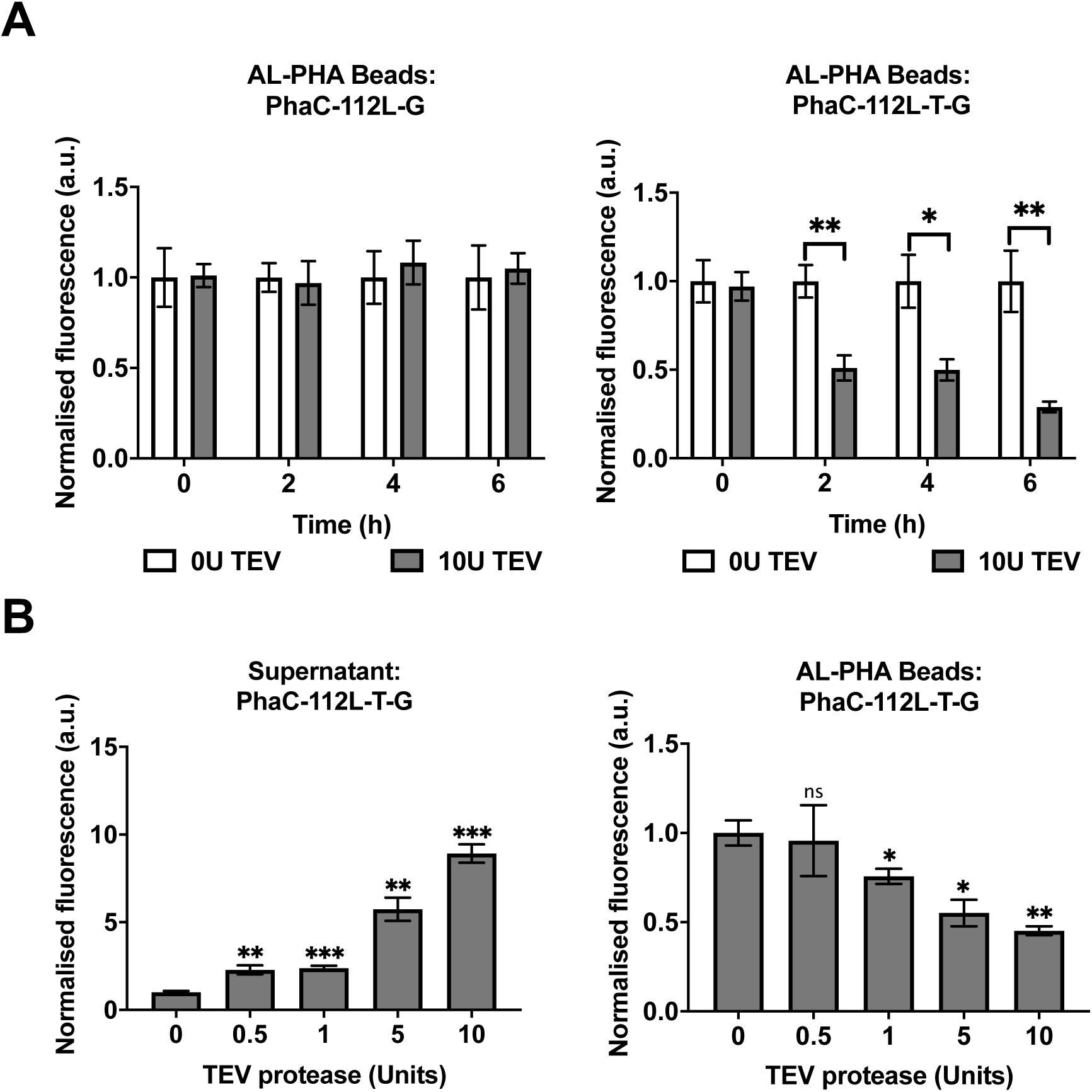
AL-PHA biosensor assay optimisation. **(A)** Time course assay. Control (PhaC-112L-G) and TEV (PhaC-112L-T-G) AL-PHA biosensor beads were incubated with 0 (0 U) or 10 units (10 U) of AcTEV protease for 0-6 hours. AL-PHA beads were analysed using flow cytometry and AL-PHA bead geometric mean (BL1-A, 488nm/530-30nm) of TEV treated beads were normalised against untreated controls of the same biosensor batch. **(B)** AL-PHA biosensor sensitivity. AL-PHA TEV biosensors were incubated for 2 hours with 0-10 units of AcTEV protease. Supernatant fluorescence data (483-14 nm/530-30nm) were normalised against untreated controls of the same biosensor batch. AL-PHA beads were analysed using flow cytometry and the geometric mean of TEV treated AL-PHA biosensor beads were normalised against untreated controls of the same biosensor batch. Error bars denote standard error of the mean, n=3-4 (AL-PHA batches), Student *t*-test *P<0.05, **P<0.01, ***P<0.001 or not statistically significant (ns).

### Generating a panel of AL-PHA protease sensitive beads to detect specific proteases

A panel of 112L AL-PHA protease biosensors were then designed and built to detect cercarial elastase or metalloproteinases, including select MMPs, ADAMs and ADAMTSs (Supplementary Table 3). Several batches of these different AL-PHA beads were generated in engineered *E. coli* and purified as described above. AL-PHA bead sizes were characterised using dynamic light scattering (DLS) and were typically 1.1 ±0.02 μm in diameter (Supplementary Fig. 8A). Flow cytometry was also used to confirm that that these AL-PHA bead biosensors were fluorescent. Essentially, 112L AL-PHA beads were typically ∼30-48 fold more fluorescent than the non-functionalised control PHAs beads (C104), which indicates correct functional assembly and surface localisation of the PhaC-fusion protein *in vivo*. (Supplementary Fig. 8B). In contrast, the MMP9 and elastase specific AL-PHA beads were only ∼4 fold more fluorescent than control beads suggesting that certain protease recognition motifs (cleavage sites) within the linker region appear to affect the correct assembly of the PhaC-sfGFP fusion protein on the beads. However, flow cytometry data showed that all 112L AL-PHA beads were functionalised and suitable for application testing despite the variability described above.

### *Schistosoma mansoni* cercarial elastase detection

The neglected tropical disease schistosomiasis is of increasing burden to global health, with estimates suggesting that 779 million people are at risk of infection, leading to an annual mortality upwards of 280,000 people in sub-Saharan Africa alone [13,14,33]. Therefore, the ability to detect this parasite is of great importance. Indeed, several trap systems have been proposed that can capture and concentrate schistosoma cercariae ready for downstream sample processing and testing [23]. To investigate whether the AL-PHA assay is applicable to the detection of *S. mansoni* – one of the principal causative agents of human schistosomiasis, we engineered AL-PHA beads specific for this parasite. Previously, we targeted the *S. mansoni* cercarial elastase activity as our marker for detection using WCB’s [22]. The cercariae utilise this elastase activity to penetrate the skin barrier, thereby enabling invasion and infection of their definitive hosts, in this case humans [16,17].

We first replaced the TEV protease recognition motif with that of *S. mansoni* cercarial elastase via inverted PCR. The elastase specific recognition motif (-SWPL-) used here and in our previous study [22], was identified using positional scanning and synthetic combinatorial library screening [17]. To test whether our 112L elastase specific AL-PHA design could detect elastase, we tested three biologically distinct *S. mansoni*-derived extracts containing soluble cercarial antigens, termed cercarial transformation fluid (SmCTF; Fig. 3A) [34]. These samples were obtained by mechanically transforming cercariae released from the intermediate snail host, *Biomphalaria glabrata* (Fig. 3A) [34]. When the elastase specific AL-PHA beads were exposed to the SmCTF samples, the beads detected elastase in SmCTF samples 2 and 3, shown by an increase in supernatant sfGFP fluorescence and a reduction in sfGFP bead fluorescence (Fig. 3B). Although SmCTF1 caused a slight reduction in bead fluorescence, no corresponding increase in supernatant fluorescence was detectable (Fig. 3B), suggesting that this sample has lower amounts of elastase. We did observe some off-target cleavage for the three SmCTF samples against the TEV protease control beads (Fig. 3B), which is likely due to non-specific proteases within the cercarial gland extract [22]. However, for two of the SmCTF samples, the elastase specific AL-PHA beads are cleaved significantly more compared to controls (Fig. 3B). This demonstrates the ability of AL-PHA beads and the appropriate controls to identify samples significantly enriched with *S. mansoni* cercarial elastase.

**Figure 3.**
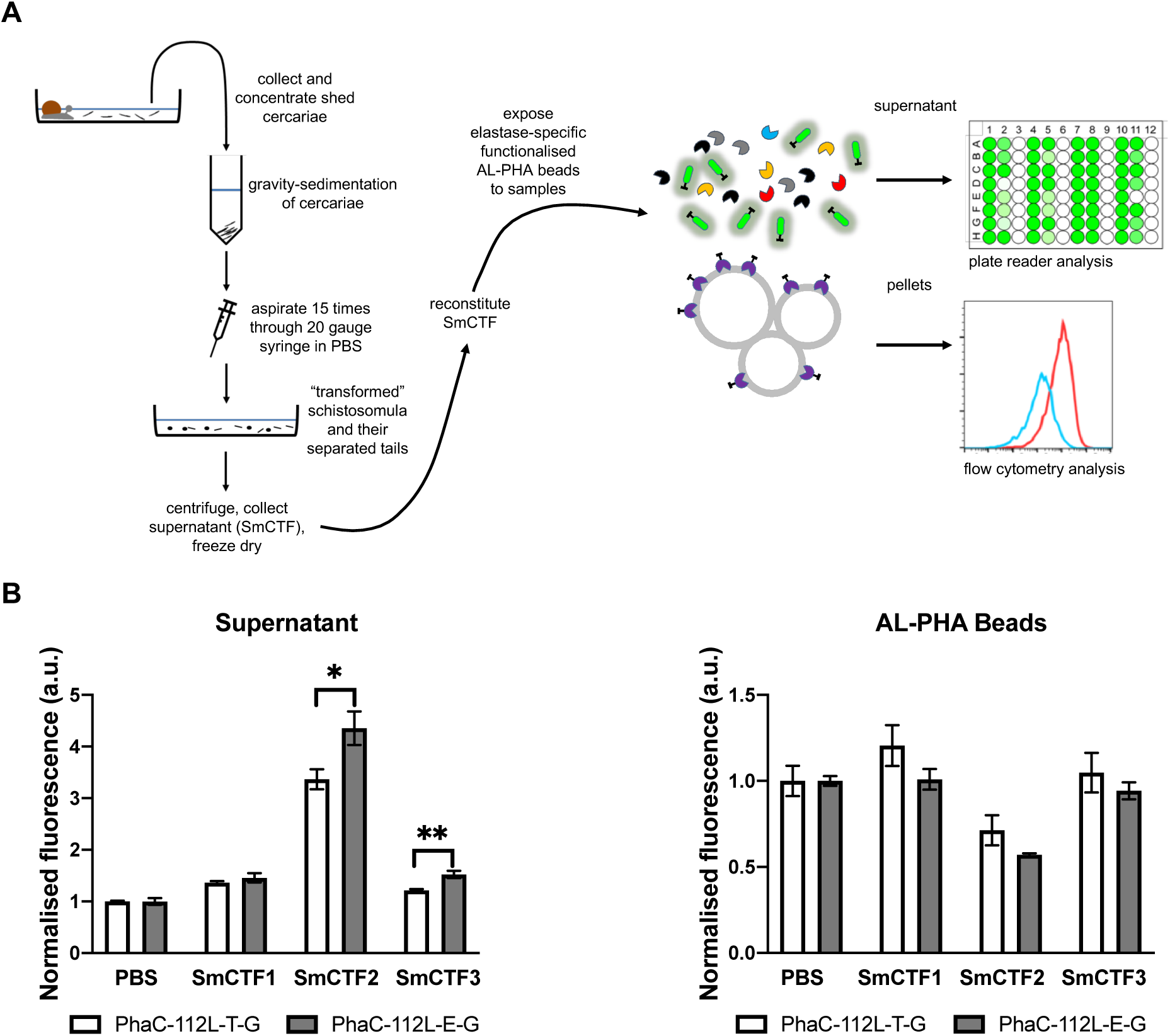
Detection of cercarial elastase in *Schistosoma mansoni* cercarial samples. **(A)** Cercarial elastase AL-PHA biosensor assay. *Schistosoma mansoni* cercariae were shed from infected snails, mechanically processed to produce *S. mansoni* cercarial transformation fluid (SmCTF) samples and then lyophilised in PBS (1X). Lyophilised SmCTF samples were reconstituted in water for AL-PHA biosensor assays. **(B)** AL-PHA TEV (PhaC-112L-T-G) and elastase (PhaC-112L-E-G) biosensors were treated with either PBS or SmCTF samples. Supernatant fluorescence data (483-14 nm/530-30nm) of SmCTF treated AL-PHA biosensors were normalised against mock-treated controls (PBS) of the same biosensor batch. AL-PHA beads were analysed using flow cytometry and the geometric mean (BL1-A, 488nm/530-30nm) of SmCTF treated AL-PHA biosensor beads were normalised against PBS controls of the same biosensor batch. Error bars denote standard error of the mean, n=3 (AL-PHA batches), Student *t*-test *P<0.05 and **P<0.01.

### Applying AL-PHA beads to detect matrix metalloproteinase MMP14

The matrix metalloproteinases (MMPs) are a family of proteases that have important physiological roles in extracellular matrix (ECM) turn-over, tissue homeostasis, immunomodulation, and cell signalling [18]. Interestingly, differential MMP14 (also called MT1-MMP) expression, protein levels or proteolytic activities are implicated in several diseases including neurodegenerative disorders and several cancers (e.g. breast, gastric, lung and ovarian cancers) [35–37]. Therefore, the detection of MMP14 proteolytic activity might be highly informative as a prognostic or diagnostic biomarker for several different biomedical applications.

To examine this, we tested MMP14-specific 112L AL-PHA beads using pre-activated recombinant MMP14 (Fig. 4). Control and MMP14-specific AL-PHA beads were treated with 0 μg or 0.55 μg (equivalent to 0 or 0.033 mU) of activated MMP14 and assayed at 37°C for 4 hours (Fig. 4A). Supernatant fluorescence levels increased by ∼2.5 fold for MMP14 treated AL-PHA beads compared to untreated controls (Fig. 4B), indicating positive detection of MMP14 proteolytic activity. For MMP14 treated AL-PHA control beads, relative supernatant fluorescence levels increased slightly (∼0.7-fold) (Fig. 4B), suggesting a low level of non-specific cleavage. Preliminary bioinformatics analysis highlighted a candidate MMP14 cleavage site within the sfGFP sequence (TGVVP-|-ILVEL; amino acids 9-18) which may account for these potential off-target affects. However, by measuring supernatant fluorescent as well as residual AL-PHA bead fluorescence, we were able to mitigate any off-target effects (Fig. 4B). Thus, our assays showed that MMP14-specific AL-PHA beads detected 0.033 mU (equivalent to 5.5 μg/ml; ∼177 nanomolar) of recombinant MMP14 activity. Since MMP14 levels within breast cancer patient serum samples (∼0.017 ±0.006 μg/ml; ∼0.3 nanomolar) [38], or plasma samples taken from liver transplant recipients, receiving a familial amyloidotic polyneuropathy liver (∼1.8 μg/ml; ∼32 nanomolar) [35], are expected to be below the observed AL-PHA bead detection limit, appropriate patient sample processing steps (e.g. protein concentration or exosome isolation protocols) will be required to concentrate MMP14 to within detectable AL-PHA biosensor levels. To test this concept, we next tested AL-PHA beads against EV-associated metalloproteinases.

**Figure 4.**
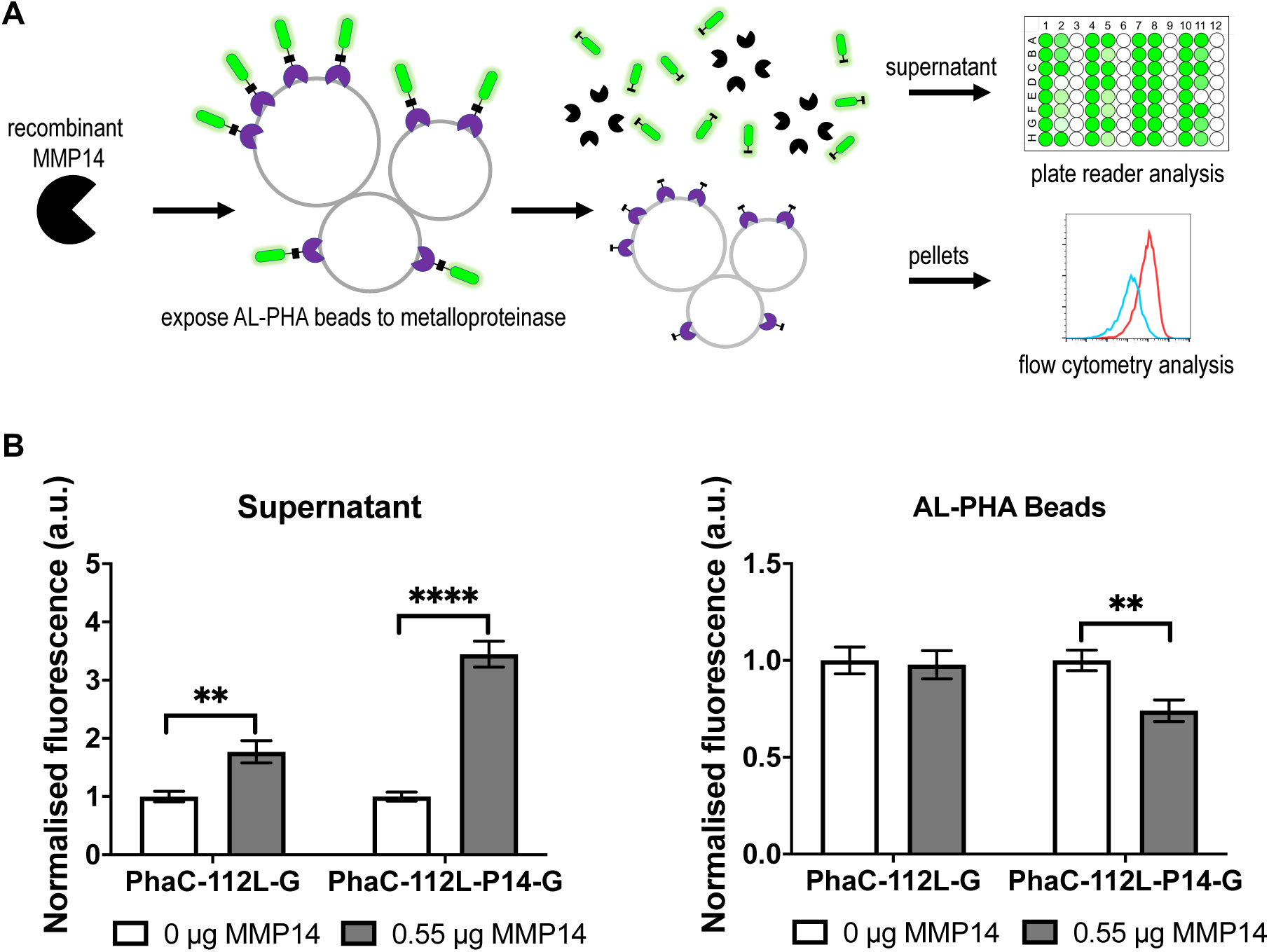
Detection of recombinant MMP14. **(A)** Matrix metalloproteinase 14 (MMP14) AL-PHA biosensor assay. **(B)** AL-PHA MMP14 (PhaC-112L-P14-G) and control (PhaC-112L-G) biosensors were treated with 0 μg or 0.55 μg of activated, recombinant MMP14. Supernatant fluorescence data (483-14 nm/530-30nm) of MMP14 treated AL-PHA biosensors were normalised against untreated controls of the same biosensor batch. AL-PHA beads were analysed using flow cytometry and the geometric mean (BL1-A, 488nm/530-30nm) of MMP14 treated AL-PHA biosensor beads were normalised against untreated controls of the same biosensor batch. Error bars denote standard error of the mean, n=8 (4 AL-PHA batches tested in duplicate), Student *t*-test **P<0.01 and ****P<0.0001.

### Detecting membrane associated metalloproteinases in extracellular vesicles using AL-PHA beads

A number of physiologically important MMPs are membrane associated (e.g. MT-MMPs and ADAMs) and one potential strategy for detection would be through the isolation of extracellular vesicles (EVs). EVs, including exosomes, are readily accessible from patient liquid biopsies and recent studies indicate that EV-associated metalloproteinases have complex roles in cancer metastasis [39,40], including lung cancer where early detection could positively impact patient outcomes [41]. Indeed, exosomes isolated from *in vitro* models of lung cancer (e.g. A549 cells) or the blood of nsclc patients exhibit increased ADAM10 activity [37,42]. For A549 cells, ADAM10 is strongly expressed and secreted within A549 EVs [42]. Therefore, ADAM10 specific AL-PHA beads may be a useful tool for lung cancer biomarker research.

After constructing ADAM10-specific 112L AL-PHA beads, we tested their ability to detect for ADAM10 proteolytic activity from purified A549 EVs (Fig. 5). Prior to the AL-PHA bead assays, A549 EVs were characterised using nanoparticle tracking analyses (EV mode diameter 88.8 ±4.8 nm; Supplementary Fig. 9), an Exo-Check dot blot array (CD81, CD63, ICAM, ANXA5, TSG101 positive; Supplementary Fig. 10), and flow cytometry surface marker characterisation (CD9, CD63 and CD81 positive; Supplementary Fig. 11). Control and ADAM10-specific AL-PHA beads were treated with either 0 μg or 50 μg (total protein) of A549 EVs using an EDTA-free protease inhibitor cocktail to inhibit a broad array of serine and cysteine proteases, but not metalloproteinases (see materials and methods). ADAM10 AL-PHA and control assays were incubated at 37°C for 4 h and then analysed, post-assay, using a plate reader and flow cytometry (Fig. 5A). Relative supernatant fluorescence levels increased by ∼0.5-fold (∼46% increase) for A549 EV-treated samples compared against untreated control samples (Fig. 5B). Furthermore, A549 EV treatment also caused a significant decrease in AL-PHA bead fluorescence (∼16% decrease), whilst control beads and supernatant fluorescence levels were unaffected (Fig. 5B). Taken together, our data show a positive detection of ADAM10 proteolytic activity from A549 EVs (50 μg total protein) which is in agreement with a previous study using lysed nsclc-patient EVs (60 μg – total protein of exosome lysates) [37]. However, in comparison to the aforementioned study [37], we envision that AL-PHA assays may enable a more simplified approach for detection. Future studies will be required to investigate if ADAM10 AL-PHA bead assays are sensitive enough to detect EV-associated ADAM10 within liquid biopsy samples or isolated patient exosomes.

**Figure 5.**
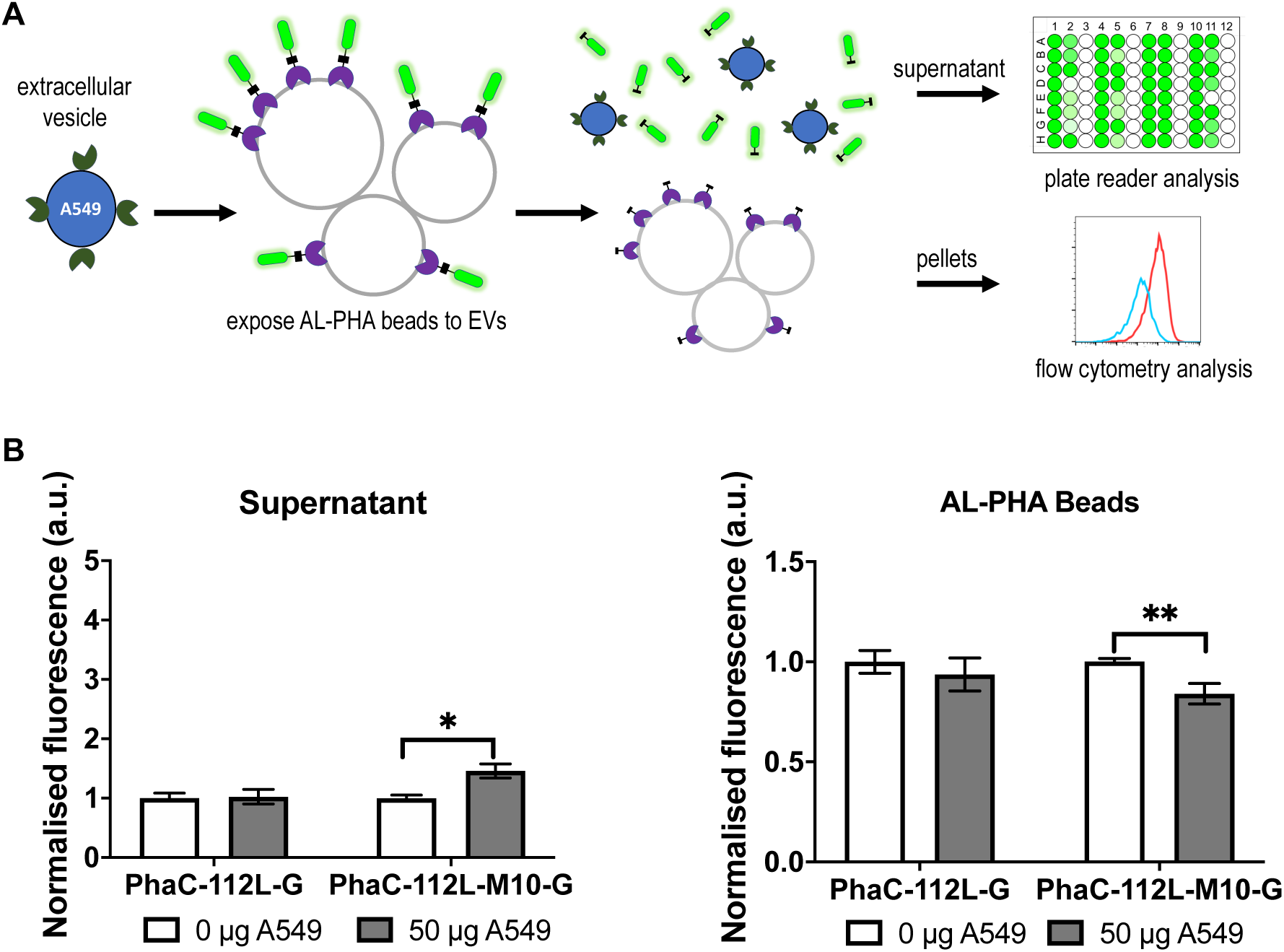
Detection of ADAM10 in A549 extracellular vesicles. **(A)** Extracellular vesicle (EV) AL-PHA biosensor assay. **(B)** AL-PHA biosensors were treated with 0 μg or 50 μg (total protein) of A549, non-small cell lung cancer cell line, extracellular vesicles. Supernatant fluorescence data (483-14 nm/530-30nm) of A549 EV treated AL-PHA biosensors were normalised against untreated controls of the same biosensor batch. AL-PHA beads were analysed using flow cytometry and the geometric mean (BL1-A, 488nm/530-30nm) of A549 EV treated AL-PHA biosensor beads were normalised against untreated controls of the same biosensor batch. Error bars denote standard error of the mean, n=4 (AL-PHA batches), Student *t*-test *P<0.05 and **P<0.01.

To support these future studies and in order to optimise AL-PHA bead screening efficiency, we devised and tested a simple, semi high-throughput assay for screening AL-PHA metalloproteinase-specific bead libraries (Fig. 6A; Supplementary Table 5). A panel of AL-PHA metalloproteinase-specific biosensors were assayed within a 96-well plate format, using conditioned media samples from HEK293 cells. These HEK293 cells were cultured at a high cell density within a hollow fibre bioreactor, to obtain concentrated cell cultures of up to ∼10^9^ cells and their secretomes (>20 kDa proteins and EVs) within ∼20 ml harvest volumes. HEK293 cells have been shown to secrete MMPs, ADAMs and ADAMTSs [43–45]. Therefore, as expected, this high-throughput screening approach enabled the successful detection of the proteolytic activities of several metalloproteinases (Fig. 6B; Supplementary Table 5). We thus, envision that this screening approach could be used in future studies to optimise AL-PHA biosensor designs or to screen for a panel of metalloproteinase activities in other biological samples.

**Figure 6.**
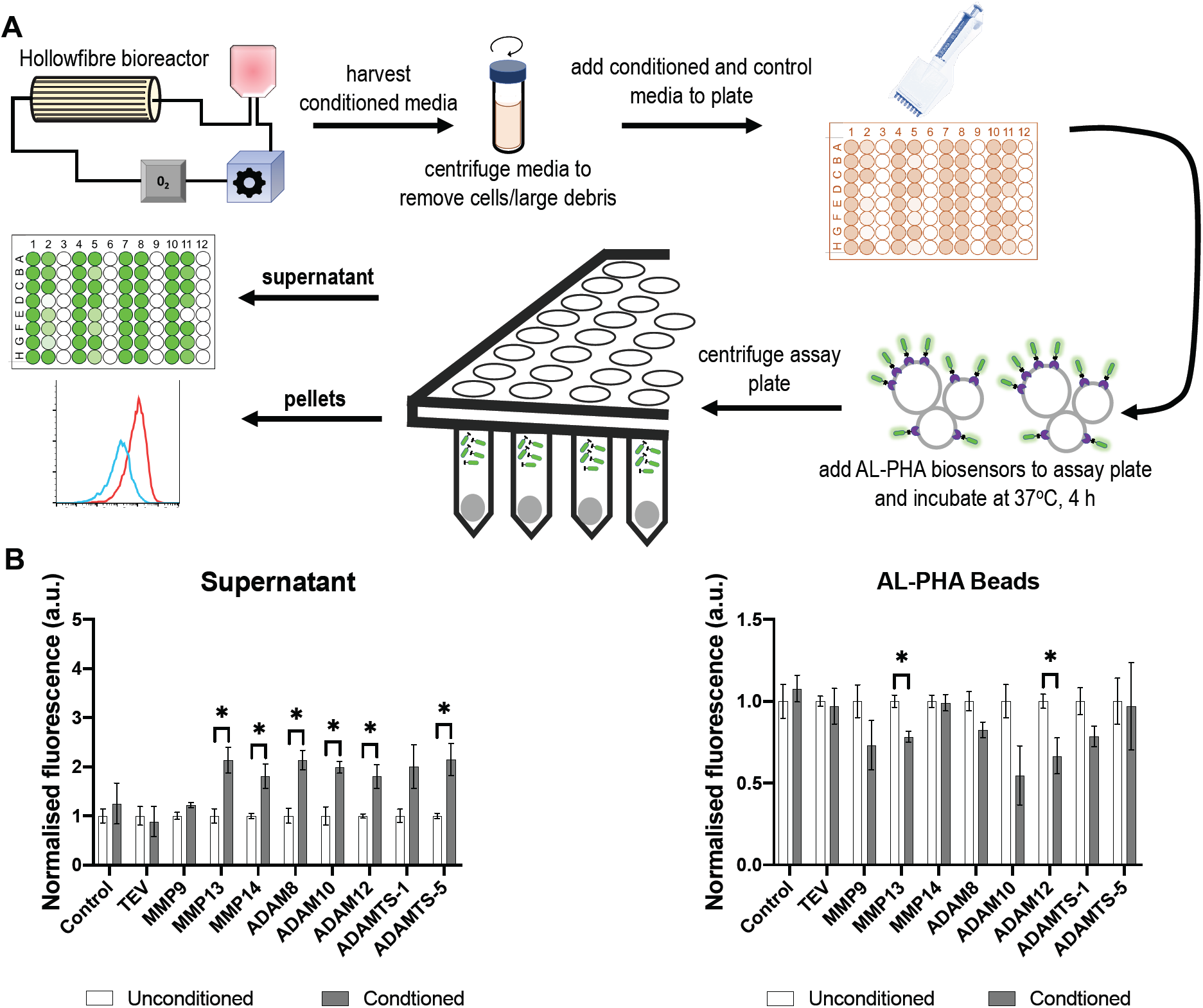
AL-PHA biosensor screening assay. **(A)** HEK293 cells were cultured to a high cell density in a hollow fibre bioreactor. Cell conditioned media were assayed in 96-well plates against a panel of control and metalloproteinase AL-PHA biosensors. **(B)** AL-PHA biosensors were treated with either conditioned or unconditioned cell culture media. Supernatant fluorescence data (483-14 nm/530-30nm) of conditioned media treated AL-PHA biosensors were normalised against unconditioned media controls of the same biosensor batch. AL-PHA beads were analysed using flow cytometry and the geometric mean (BL1-A, 488nm/530-30nm) of conditioned media treated AL-PHA biosensor beads were normalised against unconditioned media controls of the same biosensor batch. Error bars denote standard error of the mean, n=3 (AL-PHA batches), Student *t*-test *P<0.05.

## Discussion

Protease structure-function studies continue to highlight the biomedical importance of proteases in an array of communicable and non-communicable diseases [6]. Therefore, novel strategies for detecting proteolytic activity are desirable. Classical protease activity assays typically incorporate fluorogenic small molecule, peptide or nanoparticle substrates, Fluorescence Resonance Energy Transfer (FRET)-probes, electrochemical components or zymography methods [46–49]. Whereas, recent synthetic biology approaches have led to the development of more sophisticated modelling-led design strategies, the embedding of protease biosensors within smart materials and also increasingly complex whole-cell bioreporters [4,5,19,22,23]. These classical and synthetic biology approaches each have their own advantages and limitations, several of which we briefly summarise and compare alongside AL-PHA beads (Supplementary Table 6). Protease biosensor implementations, especially those intended for field or point-of-care use, must also consider responsible research and innovation requirements (e.g. implementation costs, political, regulatory and societal contexts) [23]. Beneficially, AL-PHA beads are non-living and are biodegradable [4,31] which further extends the flexibility of their implementation and safe disposal. Moreover, their modular design, ability to be measured using different approaches (visual, plate reader and flow cytometry) and capacity for use within high-throughput screening workflows, greatly extends their usefulness in terms of the different contexts where AL-PHA biosensors could potentially be implemented in the future (e.g. diagnostic labs or in the field/point-of-care). We also estimate that the raw material cost to microbially produce enough AL-PHA beads for a typical AL-PHA assay, is around 2-4 pence (GBP). Even when sample processing costs, labour and other miscellaneous costs are taken into account, AL-PHA assays are likely to remain cost competitive against competing technologies. In conclusion, we have developed a protease-detection assay using AL-PHA beads and have shown that a library of low-cost, biodegradable, AL-PHA bead protease biosensors can be utilised to detect specific protease activities. We suggest that AL-PHA beads could be a platform for implementation as low-cost global health biosensors for use in resource-limited settings.

## Materials and methods

### Bacterial strains and general growth conditions

Plasmid constructs and strains used in this study are listed in Supplementary Table 3. *E. coli* JM109 was used for both cloning and production of AL-PHA protease biosensors. For plasmid recovery *E. coli* strains were grown in Luria-Bertani (LB) medium supplemented with 34 μg/ml Chloramphenicol (final concentration) and cultured at 37°C with shaking (220 rpm). During PHAs and AL-PHA bead production *E. coli* strains were grown in Terrific-Broth (TB) supplemented with 34 μg/ml Chloramphenicol (final concentration) and 3% glucose (w/v), cultured at 37°C with shaking (220 rpm).

### Construct assembly

Empty vector plasmid EV104 was originally sourced from the 2013 distribution of the iGEM Registry of Standard Biological Parts (partsregistry.org; BBa_K608002) and was transformed during this study into *E. coli* JM109 to create strain EV104-JM109. Plasmid C104-JM109, encoding a constitutively expressed, engineered *phaCAB* operon was originally derived from our previous study [31] and was transformed during this study into *E. coli* JM109 to create strain C104-JM109. C104-JM109 was used to create wild-type, non-functionalised PHAs beads.

For 12L AL-PHA biosensor construction, the PhaC fusion (12L-G) region was ordered as a dsDNA gene synthesis fragment (GeneArt String, ThermoFisher, USA) and primer pair RK028/RK029 were used to amplify and linearise plasmid C104-JM109. In-Fusion cloning (Takara Bio, USA) of linearised C104-JM109 with the GeneArt String and transformation into *E. coli* JM109 was used to clone the 12L AL-PHA biosensor control plasmid (PhaC-12L-G; pYZW1). In order to incorporate protease recognition motifs (TEV, MMP, ADAM and ADAMTS) into the linker region, plasmid pYZW1 was used as a PCR template with appropriate primer pairs (YZW11-YZW78). The resultant PCR products were digested with *Dpn*I, phosphorylated, re-ligated and transformed into *E. coli* JM109, resulting in plasmids/strains pYZW2-pYZW29.

For 22L AL-PHA biosensor construction, the longer flexible amino acid linker (22L) region was incorporated using PCR. Briefly, 12L AL-PHA biosensor plasmids were used as templates in PCR reactions, along with primer pair YZW81/YZW82. The resultant PCR products were digested with *Dpn*I, phosphorylated, re-ligated and transformed into *E. coli* JM109, resulting in the 22L biosensor plasmids/strains PhaC-22L-G (pYZW33) and PhaC-22L-T-G (pYZW34).

For 112L AL-PHA biosensor construction, the longer flexible amino acid linker (112L) region was ordered as a dsDNA gene synthesis fragment (gBlock, IDT, USA) and primer pair YZW83/YZW84 was used to amplify and linearise all 12L AL-PHA biosensor plasmids. In-Fusion cloning (Takara Bio, USA) of linearised 12L AL-PHA biosensor plasmids with the gBlock and subsequent transformation into *E. coli* JM109 resulted in the 112L AL-PHA biosensor constructs/strains (pYZW39-pYZW48). The 112L Elastase biosensor (PhaC-112L-E-G) was cloned separately. Briefly, plasmid pYZW40 (PhaC-112L-T-G) was used as a PCR template, along with primer pair AJW671/AJW672. The resultant PCR products were digested with *Dpn*I, phosphorylated, re-ligated and transformed into *E. coli* JM109, resulting in 112L Elastase biosensor plasmid/strain PhaC-112L-E-G (pAJW290).

Oligonucleotide primers used for generating biosensor constructs and DNA sequencing are shown in Supplementary Table 4.

### Production, and characterisation of AL-PHA biosensor beads

Glycerol stocks of control (C104-JM109) or AL-PHA biosensor strains were used to inoculate 0.5 L flasks containing 100 ml Terrific Broth (TB), supplemented with 3% glucose (w/v) and 34 μg/ml Chloramphenicol (Cam). These production cultures were incubated at 37°C with shaking at 200 rpm for 24 h. Subsequently, these production cultures were harvested via centrifugation at 3220 *g* for 10 minutes at 4°C. The resultant cell pellets were washed twice in phosphate-buffered saline (PBS) (1X), before being re-suspended into 1 ml PBS (1X) per gram of the cell pellet and transferred into 2 ml microtubes. To release PHAs/AL-PHA biosensor beads, samples were sonicated using a Vibra-cell VCX130 sonicator (SONICS, Newtown, USA), with a 6 mm diameter probe, on ice (3 × 40 s with 59 s cooling interval; output frequency: 20 kHz; amplitude: 50 %). Post-lysis, samples were centrifuged 6000 *g* for 10 minutes at 4°C and then gently re-suspended (5 seconds at setting 5) using a vortex machine (Heidolph, REAX 2000). Samples were sonicated a second time (2 × 20 s with 59 s cooling interval; output frequency: 20 kHz; amplitude: 50 %) on ice and then centrifuged 6000 *g* for 10 minutes at 4°C. The supernatant was removed and the released PHAs/AL-PHA beads were re-suspended as a 20 % slurry (pellet w/v in PBS). PHAs/AL-PHA bead batches were stored at 4°C with 2 μl kanamycin (stock concentration 25 μg/ml) per microtube batch to prevent bacterial growth. PHAs and AL-PHA biosensor beads were analysed using dynamic light scattering (DLS) on a Malvern Zetasizer Nano ZS (Malvern Instruments, Malvern, UK). Harvested PHAs/AL-PHA biosensor beads were diluted 100-fold into PBS (1X) and at least three technical replicates per sample were measured at 25°C.

### SmCTF sample preparation

The three SmCTF samples tested (SmCTF1-3) were produced by BioGlab Ltd. (Nottingham, UK) as previously described [22,34]. Freeze-dried SmCTF samples were reconstituted in sterile distilled water and stored at −20°C until required.

### MMP14 activation reactions

Recombinant human MMP-14 (MT1-MMP) metalloproteinase (ab#168081, Abcam, MA, USA) was commercially sourced and activated according to the manufacturer’s instructions. Briefly, MMP14 activation reactions were 100 μl in total and consisted of 25 μl MMP14 (5 μg), 1 μl of 50 μg TPCK-trypsin per ml activation buffer (#T1426-50MG, Sigma-Aldrich, MO, USA) and 74 μl of activation buffer (50mM TRIS-HCl, pH 7.5, 150 mM NaCl, 5 mM CaCl_2_). Activation reactions were incubated at 25°C for 12 min. Post-incubation, 1 μl of 1 mg/ml aprotinin (activation buffer; #Ab146286, Abcam MA, USA) was added to inactivate TPCK-trypsin.

### Hollow fibre cell culture

HEK293 cells were cultured at a high cell density using a hollow fibre bioreactor (FiberCell Systems, Inc., MD, USA), which was configured with a 20 kDa molecular weight cut off cartridge (#C2011, FiberCell Systems, Inc., MD, USA) that essentially concentrated HEK293 cells (up to ∼10^9^ cells) and their secretomes (>20 kDa proteins and EVs) within ∼20 ml cartridge/harvest volumes. The hollow fibre cartridge was sequentially primed, pre-culture by PBS (1X; #14190144, ThermoFisher Scientific, MA, USA), serum-free Fluorobrite DMEM (#A1896702, ThermoFisher Scientific, MA, USA) and Fluorobrite DMEM (#A1896702, ThermoFisher Scientific, MA, USA) supplemented with 10% Exosome-Depleted Fetal Bovine Serum (FBS, #A25904DG, ThermoFisher Scientific, MA, USA), each for 24 h. Once primed, 1×10^8^ adherent HEK293 cells were seeded into the cartridge. HEK293 cell line validation was carried out by Eurofins (Supplementary Fig. 12). Cell growth rate was indirectly evaluated through daily monitoring of the glucose level of the cell culture medium (#GC001000, FiberCell Systems, Inc., MD, USA). The cell culture medium was changed once glucose levels significantly decreased to less than half the original level. Once glucose consumption levels significantly increased (e.g. 50% glucose consumed within 24 h) the cell culture media was changed to Fluorobrite DMEM (#A1896702, ThermoFisher Scientific, MA, USA) supplemented with 10% Chemically Defined Medium for High Density Cell Culture (#CDM HD, FiberCell Systems, Inc., MD, USA), a protein-free FBS replacement. Samples (∼ 20 ml volume) were harvested from the hollow fibre cartridge each day and HEK293 cells were cultured for a week, using serum free media prior to harvesting conditioned media for AL-PHA assays.

### AL-PHA biosensor assays

AL-PHA biosensor reactions were setup according to the optimal activity requirements of the proteases being detected. Trypsin AL-PHA assays: 50 μl (total volume) reactions were setup within 1.5ml microtubes and included the following components: 1 μl of purified AL-PHA beads (PhaC-22L-G), 0 or 1 μg Sequencing Grade Modified Trypsin (0 or 10 μl; #V5111, Promega, WI, USA) that was reconstituted within trypsin re-suspension buffer (50 mM acetic acid; #V542A, Promega, WI, USA), and 39 μl or 49 μl 250 mM Tris buffer. These reactions were incubated for 2 h at 37°C with 220 rpm shaking (Eppendorf, Thermo Mixer C). AcTEV protease AL-PHA assays: 100 μl (total volume) reactions were setup within 1.5 ml microtubes and included the following components: 1 μl of purified AL-PHA beads, 0-10 U AcTEV (0-1 μl; #12575015, ThermoFisher Scientific, MA, USA), 1 μl of dithiothreitol (DTT) 0.1M, 5 μl of TEV reaction buffer (20X; 1M Trix-HCl pH8.0, 10mM EDTA), and 92-93 μl of PBS (1X). These reactions were incubated for 0-6 h at 30°C with 500 rpm shaking (Eppendorf, Thermo Mixer C). *S. mansoni* cercarial elastase AL-PHA assays: 100 μl (total volume) reactions were setup within 1.5 ml microtubes and included the following components, 1 μl of purified AL-PHA beads, 10 μl of reconstituted SmCTF sample and 89 μl PBS (1X). These reactions were incubated for 2 h at 30°C with 500 rpm shaking (Eppendorf, Thermo Mixer C). MMP14 AL-PHA assays: 100 μl (total volume) reactions were setup within 1.5ml microtubes and included the following components, 1 μl of purified AL-PHA beads, 0 or 0.55 μg of activated MMP14 (0 or 11 μl), 87 or 98 μl of activation buffer (50mM TRIS-HCl, pH 7.5, 150 mM NaCl, 5 mM CaCl_2_) and 1μl of 1 mg/ml aprotinin (activation buffer; #Ab146286, Abcam MA, USA). These reactions were incubated for 4 h at 37°C with 500 rpm shaking (Eppendorf, Thermo Mixer C). EV-associated ADAM10 AL-PHA assays: 100 μl (total volume) reactions were setup within 1.5 ml microtubes and included the following components, 1 μl of purified AL-PHA beads, 0 or 50 μg of A549 EVs (0 or 50 μl; #HBM-A549-100/5, HansaBioMed, Tallinn, Estonia), 4 μl of cOmplete™ EDTA-free protease inhibitor cocktail (50X; #11873580001, Sigma-Aldrich, MO, USA) and 45 μl or 95 μl PBS (1X). These reactions were incubated for 4 h at 37°C with 500 rpm shaking (Eppendorf, Thermo Mixer C). HEK293 conditioned media AL-PHA assays: 100 μl (total volume) reactions were setup within the wells of a 96-well plate (#655076, Greiner Bio-One, Kremsmünster, Austria) and included the following components, 0 or 1 μl of purified control (C104) PHAs beads, 0 μl or 1 μl of purified AL-PHA beads, 0 or 99 μl of fresh, serum-free non-conditioned media (Fluorobrite DMEM; #A1896702, ThermoFisher Scientific, MA, USA) and 0 or 99 μl of HEK293 conditioned cell media (see hollow fibre cell culture materials and methods section). These 96-well assay plates were incubated for 4 h, within a plate reader (CLARIOstar, BMG, Ortenberg, Germany) at 37°C with 300 rpm shaking.

Post-incubation, AL-PHA assay tubes were centrifuged ≥6000 *g* for 10 minutes, whilst AL-PHA assay plates were centrifuged 500 *g* for 10 minutes (MPS 1000 Mini PCR Plate Spinner, Sigma-Aldrich, MO, USA) in order to pellet AL-PHA beads. Post-centrifugation, 20 μl of each AL-PHA assay supernatant were separately sampled and transferred to a black, μClear 384 well plate (#781096, Greiner Bio-One, Kremsmünster, Austria) for plate reader analyses (Excitation 483-14 nm/Emission 530-30; CLARIOstar, BMG, Ortenberg, Germany). Supernatant fluorescence measurements of protease treated AL-PHA biosensors were normalised against non-treated controls of the same biosensor batch. Whilst, pelleted AL-PHA beads were re-suspended into 1 ml PBS (1X) and around 10,000 events for each sample were analysed using flow cytometry (Attune NxT, ThermoFisher Scientific, MA, USA). The geometric mean (BL1-A, Excitation 488 nm/Emission 530/30 nm) of protease treated AL-PHA beads were normalised against non-treated controls of the same biosensor batch and at least three batches of each AL-PHA biosensor were tested in all AL-PHA assays. The flow cytometry gating strategy used (Supplementary Fig. 4) was also validated with the assistance of 1 μm bead standards (#F13839, Thermo Fisher Scientific, MA, USA).

### GFP Calibration curve

Super folder green fluorescent protein (sfGFP) samples were purified from AL-PHA beads and used to create an AL-PHA GFP calibration curve (Supplementary Fig. 5). Briefly, four 1.5 ml microtubes were setup with 4 μl of purified AL-PHA beads (PhaC-112L-T-G) and 96 μl PBS (1X). These samples were centrifuged (6000 *g* for 10 minutes) to wash and pellet the AL-PHA beads and post-centrifugation the supernatant was carefully removed. AL-PHA beads were then re-suspended within 100 μl (total volume) of AcTEV reaction solutions that included the following components: 4 μl of re-suspended AL-PHA beads (PhaC-112L-T-G), 20 U AcTEV (2 μl; #12575015, ThermoFisher Scientific, MA, USA), 1 μl of dithiothreitol (DTT) 0.1M, 5 μl of TEV reaction buffer (20X; 1M Trix-HCl pH8.0, 10mM EDTA), and 88 μl of PBS (1X). These reactions were incubated for 3 h at 30°C with 500 rpm shaking (Thermo Mixer C, Eppendorf, Germany). Post-incubation, these reactions were centrifuged 6000 *g* for 10 minutes to pellet the AL-PHA beads. Supernatants containing proteolytically released sfGFP were combined and concentrated at 45°C for 20 min with VAQ setting, using an Eppendorf concentrator (Eppendorf, Germany). The protein concentration of purified sfGFP samples was determined, according to the manufactures’ guidance, using a Qubit 3 Fluorometer (Thermo Fisher Scientific, MA, USA) and a Qubit Protein Assay Kit (#Q33211, Thermo Fisher Scientific, MA, USA). The sfGFP purification samples were combined and analysed via sodium dodecyl sulphate (SDS)-polyacrylamide gel electrophoresis (PAGE) using 4–12% Bis-Tris gels (NuPAGE Novex, Thermo Fisher Scientific, MA, USA), followed by western blot analysis using a HRP-conjugated GFP-specific polyclonal antibody (1:4,000 dilution; #A10260, Thermo Fisher Scientific, MA, USA). Western blots were developed by enhanced chemiluminence (ECL). Additionally, the combined sfGFP purification sample was used to setup a GFP calibration curve. Calibration samples for each sfGFP concentration were setup in quadruplicate then aliquoted into 384-well plates (#781096, Greiner Bio-One, Kremsmünster, Austria) and measured using a CLARIOstar plate reader (Excitation 483-14 nm/Emission 530-30; BMG, Ortenberg, Germany).

### Extracellular vesicle characterisation

Nanoparticle tracking analysis (NTA): 1 μL A549 EVs (1 mg/ml; #HBM-A549-100/5, HansaBioMed, Tallinn, Estonia) were diluted (1:1000) within particle-free DPBS (1X; #14190144, Thermo Fisher Scientific), gently pipetted for 10 s and then aliquoted into a deep-well, 96-well plate. Samples from the 96-well plate were injected into a NanoSight NS300 NTA instrument equipped with a NanoSight Sample Assistant (Malvern Instruments, UK) autosampler system. The NTA images were recorded and analysed to obtain the concentration and distribution of the sample particles. Additional software version and measurement settings are shown in supplementary figure 9. Exo-Check Array: A549 protein markers were characterised using a commercially sourced Exo-Check Exosome Antibody Array kit (#EXORAY200A-4, System Biosciences, CA, USA). A549 EVs (100 μg; #HBM-A549-100/5, HansaBioMed, Tallinn, Estonia) were lysed and processed according to the manufacturer’s guidelines and the developed dot blot array was imaged using a ChemiDoc imaging system (Bio-Rad Laboratories Inc., USA) (Supplementary Fig. 10). Flow cytometry analysis of EV surface markers: Several A549 EV samples were setup and included the following components: 25 μl of A549 EVs (25 μg; #HBM-A549-100/5, HansaBioMed, Tallinn, Estonia) were incubated with 20 μl of Exosome-Human CD63 Dynabeads (#10606D, Thermo Fisher Scientific, MA, USA) and 53 μl of DPBS (1X; #14190144, Thermo Fisher Scientific) at 25°C for 1 hour with shaking 1000 rpm (Thermo Mixer C, Eppendorf, Germany) in order to bind CD63+ EVs. Post-incubation, samples were transferred to a magnetic rack and unbound EVs were washed away using DPBS (1X). Dynabead-EV samples were re-suspended into 98 μl DPBS and 2 μl of one of the following antibodies: IgG1-PE (#130-113-200, Miltenyi Biotec, Germany), CD9-PE, human (#130-103-955, Miltenyi Biotec, Germany), CD63-PE, human (#130-100-153, Miltenyi Biotec, Germany) or CD81-PE, human (#130-118-342, Miltenyi Biotec, Germany). Samples were subsequently incubated at 25°C for 1 hour with shaking 1000 rpm (Thermo Mixer C, Eppendorf, Germany) in order to fluorescently label EVs. Post-incubation, samples were transferred to a magnetic rack and unbound antibodies were washed away using DPBS (1X). Dynabead-EV samples were then re-suspended into 500 μl DPBS and analysed using flow cytometry (Attune NxT, ThermoFisher Scientific, MA, USA; YL1-A Excitation 561 nm/Emission 585-16 nm).

### Statistics

Statistical analysis (standard error of the mean (s.e.m.) and unpaired *t*-test) was carried out on at least three experimental replicates using GraphPad Prism 8.4.2 (GraphPad Software Inc., La Jolla, California) and flow cytometry data analysis was performed using FlowJo (vX 10.5.3) software.

## Acknowledgements

Dr Richard Kelwick is supported by the Cancer Research UK Imperial Centre Development Fund and until recently, was supported by a BBSRC-funded Royal Society of Edinburgh Enterprise Fellowship. Dr Alexander Webb, Dr Fiona Allan, Dr Aidan Emery, Prof. Michael Templeton and Prof. Paul Freemont are supported by the UK Government’s Global Challenges Research Fund (GCRF) through the EPSRC grant [EP/P028519/1], as part of the WISER project. We also acknowledge the support of Imperial Confidence in Concept (MRC/EPSRC), Imperial College London EPSRC Impact Acceleration Account [EP/R511547/1], EPSRC grant [EP/L011573/1], and BBSRC grant [BB/L027852/1]. We thank Prof. Michael Doenhoff for SmCTF samples. We also thank Dr Matthew Haines and colleagues in the Section of Structural and Synthetic Biology, as well as Prof. Michael Seckl, Dr Roy Rajat, Dr David McClymont, and Prof. Dylan Edwards for their advice.

## Author contributions

R.K., A.W., Y.W., A.H., and P.F. conceived and designed the experiments. R.K., A.W., Y.W., and A.H. performed the experiments. R.K., A.W., Y.W., A.H., F.A., A.E., M.T., and P.F. wrote and/or edited the manuscript.

## Supplementary Data

Supplementary data associated with this article can be found in the online version.

