## Supplementary for "AL-PHA beads: bioplastic-based protease biosensors for global health applications"

^1^Section of Structural and Synthetic Biology, Department of Infectious Disease, Imperial College London, London, SW7 2AZ, UK. ^2^Department of Life Sciences, Natural History Museum, London, SW7 5BD, UK. ^3^Department of Civil and Environmental Engineering, Imperial College London, SW7 2AZ, UK. ^4^The London Biofoundry, Imperial College Translation & Innovation Hub, White City Campus, 80 Wood Lane, London W12 0BZ, UK. ^5^UK Dementia Research Institute Care Research and Technology Centre, Imperial College London, Hammersmith Campus, Du Cane Road, London, W12 0NN

^#^Joint First Authors. *To whom correspondence should be addressed:

Prof Paul Freemont:

Section of Structural and Synthetic Biology, Department of Infectious Disease, Sir Alexander Fleming Building, South Kensington Campus, Exhibition Road, London, SW7 2AZ, UK

**Supplementary Table 1. Protease cleavage recognition sites.**

| Proteases | Secreted /Membrane-bound | Protease recognition motif(s) | Reference(s) |
| --- | --- | --- | --- |
| Tobacco Etch Virus (TEV) Protease | | | |
| TEV | Secreted | ENLYFQG | (Phan et al., 2002) |
| Matrix Metalloproteinase (MMP) | | | |
| MMP1 | Secreted | PQALVAALD | (Eckhard et al., 2016) |
| MMP2 | Secreted | PGPPGPGGLNGPPGDP | (Wang et al., 2017) |
| MMP3 | Secreted | PGGPGPAGLRGPPGDP | (Wang et al., 2017) |
| MMP7 | Secreted | GGGPGPGGLGGGGGGG | (Wang et al., 2017) |
| MMP8 | Secreted | PGPPGPSGLRGPPGPP | (Wang et al., 2017) |
| MMP9 | Secreted | PGPPGPAGLRGPPGGP | (Wang et al., 2017) |
| MMP10 | Secreted | GPAGLLGP | (Schlage et al., 2014) |
| MMP11 | Secreted | AANLTR | (Pan et al., 2003) |
| MMP12 | Secreted | GGAAGPAGLGGAAGAG | (Wang et al., 2017) |
| MMP13 | Secreted | PAGLVADAD | (Eckhard et al., 2016) |
| MMP14/ MT1-MMP | Membrane Bound | PAGLVGPPN | (Eckhard et al., 2016) |
| MMP15/ MT2-MMP | Membrane Bound | MMPQGLRINV | (Kukreja et al., 2015) |
| MMP16/ MT3-MMP | Membrane Bound | MMPQGLRMRV | (Kukreja et al., 2015) |
| MMP17/ MT4-MMP | Membrane Bound | PGPEPEVLPV | (Kukreja et al., 2015) |
| MMP20 | Secreted | QAAMDMRASG | (Kukreja et al., 2015) |
| MMP24/ MT5-MMP | Membrane Bound | MMPLGMRMRR | (Kukreja et al., 2015) |
| MMP25/ MT6-MMP | Membrane Bound | PMPMEEVMPR | (Kukreja et al., 2015) |
| MMP26 | Secreted | KPISLISS | (Park et al., 2002) |
| A Disintegrin and Metalloproteinase (ADAM) | | | |
| ADAM8 | Membrane Bound | PAAAQRLR | (Rawlings et al., 2018) |
| ADAM9 | Membrane Bound | PPAASSLR | (Rawlings et al., 2018) |
| ADAM10 | Membrane Bound | SAQAVVSQ | (Rawlings et al., 2018) |
| ADAM12 | Membrane Bound | AVRSSSRTPSDK | (Rawlings et al., 2018) |
| ADAM15 | Membrane Bound | EQLRMKLP | (Rawlings et al., 2018) |
| ADAM17 | Membrane Bound | LAAAVVSS | (Rawlings et al., 2018) |
| ADAM19 | Membrane Bound | PVAASSLR | (Rawlings et al., 2018) |
| ADAM28 | Membrane Bound | SRLRAYLL | (Rawlings et al., 2018) |
| ADAM30 | Membrane Bound | PCQSASSASALGGVKVER | (Letronne et al., 2016) |
| ADAM33 | Membrane Bound | LVEALYLV | (Rawlings et al., 2018) |
| ADAMDEC1 | Membrane Bound | AFKCLKDG | (Rawlings et al., 2018) |
| A Disintegrin and Metalloproteinase with Thrombospondin Motifs (ADAMTS) | | | |
| ADAMTS1 | Secreted | VAQEARGG | (Rawlings et al., 2018) |
| ADAMTS2 | Secreted | NFAPQMSY | (Bekhouche and Colige, 2015) |
| ADAMTS4 | Secreted | LPLEFRGGTV | (Rawlings et al., 2018) |
| ADAMTS5 | Secreted | PAPEGRGGMV | (Rawlings et al., 2018) |
| ADAMTS8 | Secreted | TEGEARGS | (Rawlings et al., 2018) |
| ADAMTS9 | Secreted | TAQEAGEG | (Rawlings et al., 2018) |
| ADAMTS13 | Secreted | NLVYMVTG | (Rawlings et al., 2018) |
| ADAMTS15 | Secreted | HFQEARGWTV | (Biniossek et al., 2016) |
| ADAMTS20 | Secreted | EAAEARRG (mouse-type) | (Rawlings et al., 2018) |
| Schistosome Cercarial Elastase | |  |  |
| Elastase | Secreted | SWPL | (Salter et al., 2002) |

**Supplementary Table 2. AL-PHA biosensor fusion protein sequences.**

| **Biosensor name** | **Description** | **Fusion Protein Amino Acid Sequence** |
| --- | --- | --- |
| PhaC-12L-G | 12L control AL-PHA biosensor.  PhaC-fusion amino acid sequence shown. | **MATGKGAAASTQEGKSQPFKVTPGPFDPATWLEWSRQWQGTEGNGHAAAS**  **GIPGLDALAGVKIAPAQLGDIQQRYMKDFSALWQAMAEGKAEATGPLHDRR**  **FAGDAWRTNLPYRFAAAFYLLNARALTELADAVEADAKTRQRIRFAISQWV**  **DAMSPANFLATNPEAQRLLIESGGESLRAGVRNMMEDLTRGKISQTDESAFE**  **VGRNVAVTEGAVVFENEYFQLLQYKPLTDKVHARPLLMVPPCINKYYILDLQ**  **PESSLVRHVVEQGHTVFLVSWRNPDASMAGSTWDDYIEHAAIRAIEVARDISG**  **QDKINVLGFCVGGTIVSTALAVLAARGEHPAASVTLLTTLLDFADTGILDVFVD**  **EGHVQLREATLGGGAGAPCALLRGLELANTFSFLRPNDLVWNYVVDNYLKGN**  **TPVPFDLLFWNGDATNLPGPWYCWYLRHTYLQNELKVPGKLTVCGVPVDLAS**  **IDVPTYIYGSREDHIVPWTAAYASTALLANKLRFVLGASGHIAGVINPPAKNKRS**  **HWTNDALPESPQQWLAGAIEHHGSWWPDWTAWLAGQAGAKRAAPANYGNA**  **RYRAIEPAPGRYVKAKAVLAVIDKRGGGGGMRKGEELFTGVVPILVELDGD**  **VNGHKFSVRGEGEGDATNGKLTLKFICTTGKLPVPWPTLVTTLTYGVQCF**  **ARYPDHMKQHDFFKSAMPEGYVQERTISFKDDGTYKTRAEVKFEGDTLVN**  **RIELKGIDFKEDGNILGHKLEYNFNSHNVYITADKQKNGIKANFKIRHNVED**  **GSVQLADHYQQNTPIGDGPVLLPDNHYLSTQSVLSKDPNEKRDHMVLLEF**  **VTAAGITHGMDELYK*** |
| PhaC-22L-G | 22L control AL-PHA biosensor.  PhaC-fusion amino acid sequence shown. | **MATGKGAAASTQEGKSQPFKVTPGPFDPATWLEWSRQWQGTEGNGHAAAS**  **GIPGLDALAGVKIAPAQLGDIQQRYMKDFSALWQAMAEGKAEATGPLHDR**  **RFAGDAWRTNLPYRFAAAFYLLNARALTELADAVEADAKTRQRIRFAISQ**  **WVDAMSPANFLATNPEAQRLLIESGGESLRAGVRNMMEDLTRGKISQTDE**  **SAFEVGRNVAVTEGAVVFENEYFQLLQYKPLTDKVHARPLLMVPPCINKY**  **YILDLQPESSLVRHVVEQGHTVFLVSWRNPDASMAGSTWDDYIEHAAIRA**  **IEVARDISGQDKINVLGFCVGGTIVSTALAVLAARGEHPAASVTLLTTLL**  **DFADTGILDVFVDEGHVQLREATLGGGAGAPCALLRGLELANTFSFLRPN**  **DLVWNYVVDNYLKGNTPVPFDLLFWNGDATNLPGPWYCWYLRHTYLQNEL**  **KVPGKLTVCGVPVDLASIDVPTYIYGSREDHIVPWTAAYASTALLANKLR**  **FVLGASGHIAGVINPPAKNKRSHWTNDALPESPQQWLAGAIEHHGSWWPD**  **WTAWLAGQAGAKRAAPANYGNARYRAIEPAPGRYVKAKAGQQSGGSSQGV**  **LAVIDKRGGGGGMRKGEELFTGVVPILVELDGDVNGHKFSVRGEGEGDAT**  **NGKLTLKFICTTGKLPVPWPTLVTTLTYGVQCFARYPDHMKQHDFFKSAM**  **PEGYVQERTISFKDDGTYKTRAEVKFEGDTLVNRIELKGIDFKEDGNILG**  **HKLEYNFNSHNVYITADKQKNGIKANFKIRHNVEDGSVQLADHYQQNTPI**  **GDGPVLLPDNHYLSTQSVLSKDPNEKRDHMVLLEFVTAAGITHGMDELYK**  ***** |
| PhaC-112L-G | 112L control AL-PHA biosensor.  PhaC-fusion amino acid sequence shown. | **MATGKGAAASTQEGKSQPFKVTPGPFDPATWLEWSRQWQGTEGNGHAAAS**  **GIPGLDALAGVKIAPAQLGDIQQRYMKDFSALWQAMAEGKAEATGPLHDR**  **RFAGDAWRTNLPYRFAAAFYLLNARALTELADAVEADAKTRQRIRFAISQ**  **WVDAMSPANFLATNPEAQRLLIESGGESLRAGVRNMMEDLTRGKISQTDE**  **SAFEVGRNVAVTEGAVVFENEYFQLLQYKPLTDKVHARPLLMVPPCINKY**  **YILDLQPESSLVRHVVEQGHTVFLVSWRNPDASMAGSTWDDYIEHAAIRA**  **IEVARDISGQDKINVLGFCVGGTIVSTALAVLAARGEHPAASVTLLTTLL**  **DFADTGILDVFVDEGHVQLREATLGGGAGAPCALLRGLELANTFSFLRPN**  **DLVWNYVVDNYLKGNTPVPFDLLFWNGDATNLPGPWYCWYLRHTYLQNEL**  **KVPGKLTVCGVPVDLASIDVPTYIYGSREDHIVPWTAAYASTALLANKLR**  **FVLGASGHIAGVINPPAKNKRSHWTNDALPESPQQWLAGAIEHHGSWWPD**  **WTAWLAGQAGAKRAAPANYGNARYRAIEPAPGRYVKAKAGQQSGGSSQGG**  **SGSSSGGQGGGSQQSSSGGSQQSQQSQGSGQQSGQQGSQGQQSGGSSQGG**  **SGSSSGGQGGGSQQSSSGGSQQSQQSQGSGQQSGQQGSQVLAVIDKRGGG**  **GGMRKGEELFTGVVPILVELDGDVNGHKFSVRGEGEGDATNGKLTLKFIC**  **TTGKLPVPWPTLVTTLTYGVQCFARYPDHMKQHDFFKSAMPEGYVQERTI**  **SFKDDGTYKTRAEVKFEGDTLVNRIELKGIDFKEDGNILGHKLEYNFNSH**  **NVYITADKQKNGIKANFKIRHNVEDGSVQLADHYQQNTPIGDGPVLLPDN**  **HYLSTQSVLSKDPNEKRDHMVLLEFVTAAGITHGMDELYK*** |
| PhaC-12L-T-G | 12L TEV-specific AL-PHA biosensor.  PhaC-fusion amino acid sequence shown. | **MATGKGAAASTQEGKSQPFKVTPGPFDPATWLEWSRQWQGTEGNGHAAAS**  **GIPGLDALAGVKIAPAQLGDIQQRYMKDFSALWQAMAEGKAEATGPLHDR**  **RFAGDAWRTNLPYRFAAAFYLLNARALTELADAVEADAKTRQRIRFAISQ**  **WVDAMSPANFLATNPEAQRLLIESGGESLRAGVRNMMEDLTRGKISQTDE**  **SAFEVGRNVAVTEGAVVFENEYFQLLQYKPLTDKVHARPLLMVPPCINKY**  **YILDLQPESSLVRHVVEQGHTVFLVSWRNPDASMAGSTWDDYIEHAAIRA**  **IEVARDISGQDKINVLGFCVGGTIVSTALAVLAARGEHPAASVTLLTTLL**  **DFADTGILDVFVDEGHVQLREATLGGGAGAPCALLRGLELANTFSFLRPN**  **DLVWNYVVDNYLKGNTPVPFDLLFWNGDATNLPGPWYCWYLRHTYLQNEL**  **KVPGKLTVCGVPVDLASIDVPTYIYGSREDHIVPWTAAYASTALLANKLR**  **FVLGASGHIAGVINPPAKNKRSHWTNDALPESPQQWLAGAIEHHGSWWPD**  **WTAWLAGQAGAKRAAPANYGNARYRAIEPAPGRYVKAKAVLAVIDKRGGG**  **GENLYFQGGMRKGEELFTGVVPILVELDGDVNGHKFSVRGEGEGDATNGK**  **LTLKFICTTGKLPVPWPTLVTTLTYGVQCFARYPDHMKQHDFFKSAMPEG**  **YVQERTISFKDDGTYKTRAEVKFEGDTLVNRIELKGIDFKEDGNILGHKL**  **EYNFNSHNVYITADKQKNGIKANFKIRHNVEDGSVQLADHYQQNTPIGDG**  **PVLLPDNHYLSTQSVLSKDPNEKRDHMVLLEFVTAAGITHGMDELYK*** |
| PhaC-22L-T-G | 22L TEV-specific AL-PHA biosensor.  PhaC-fusion amino acid sequence shown. | **MATGKGAAASTQEGKSQPFKVTPGPFDPATWLEWSRQWQGTEGNGHAAAS**  **GIPGLDALAGVKIAPAQLGDIQQRYMKDFSALWQAMAEGKAEATGPLHDR**  **RFAGDAWRTNLPYRFAAAFYLLNARALTELADAVEADAKTRQRIRFAISQ**  **WVDAMSPANFLATNPEAQRLLIESGGESLRAGVRNMMEDLTRGKISQTDE**  **SAFEVGRNVAVTEGAVVFENEYFQLLQYKPLTDKVHARPLLMVPPCINKY**  **YILDLQPESSLVRHVVEQGHTVFLVSWRNPDASMAGSTWDDYIEHAAIRA**  **IEVARDISGQDKINVLGFCVGGTIVSTALAVLAARGEHPAASVTLLTTLL**  **DFADTGILDVFVDEGHVQLREATLGGGAGAPCALLRGLELANTFSFLRPN**  **DLVWNYVVDNYLKGNTPVPFDLLFWNGDATNLPGPWYCWYLRHTYLQNEL**  **KVPGKLTVCGVPVDLASIDVPTYIYGSREDHIVPWTAAYASTALLANKLR**  **FVLGASGHIAGVINPPAKNKRSHWTNDALPESPQQWLAGAIEHHGSWWPD**  **WTAWLAGQAGAKRAAPANYGNARYRAIEPAPGRYVKAKAGQQSGGSSQGV**  **LAVIDKRGGGGENLYFQGGMRKGEELFTGVVPILVELDGDVNGHKFSVRG**  **EGEGDATNGKLTLKFICTTGKLPVPWPTLVTTLTYGVQCFARYPDHMKQH**  **DFFKSAMPEGYVQERTISFKDDGTYKTRAEVKFEGDTLVNRIELKGIDFK**  **EDGNILGHKLEYNFNSHNVYITADKQKNGIKANFKIRHNVEDGSVQLADH**  **YQQNTPIGDGPVLLPDNHYLSTQSVLSKDPNEKRDHMVLLEFVTAAGITH**  **GMDELYK*** |
| PhaC-112L-T-G | 112L TEV-specific AL-PHA biosensor.  PhaC-fusion amino acid sequence shown. | **MATGKGAAASTQEGKSQPFKVTPGPFDPATWLEWSRQWQGTEGNGHAAAS**  **GIPGLDALAGVKIAPAQLGDIQQRYMKDFSALWQAMAEGKAEATGPLHDR**  **RFAGDAWRTNLPYRFAAAFYLLNARALTELADAVEADAKTRQRIRFAISQ**  **WVDAMSPANFLATNPEAQRLLIESGGESLRAGVRNMMEDLTRGKISQTDE**  **SAFEVGRNVAVTEGAVVFENEYFQLLQYKPLTDKVHARPLLMVPPCINKY**  **YILDLQPESSLVRHVVEQGHTVFLVSWRNPDASMAGSTWDDYIEHAAIRA**  **IEVARDISGQDKINVLGFCVGGTIVSTALAVLAARGEHPAASVTLLTTLL**  **DFADTGILDVFVDEGHVQLREATLGGGAGAPCALLRGLELANTFSFLRPN**  **DLVWNYVVDNYLKGNTPVPFDLLFWNGDATNLPGPWYCWYLRHTYLQNEL**  **KVPGKLTVCGVPVDLASIDVPTYIYGSREDHIVPWTAAYASTALLANKLR**  **FVLGASGHIAGVINPPAKNKRSHWTNDALPESPQQWLAGAIEHHGSWWPD**  **WTAWLAGQAGAKRAAPANYGNARYRAIEPAPGRYVKAKAGQQSGGSSQGG**  **SGSSSGGQGGGSQQSSSGGSQQSQQSQGSGQQSGQQGSQGQQSGGSSQGG**  **SGSSSGGQGGGSQQSSSGGSQQSQQSQGSGQQSGQQGSQVLAVIDKRGGG**  **GENLYFQGGMRKGEELFTGVVPILVELDGDVNGHKFSVRGEGEGDATNGK**  **LTLKFICTTGKLPVPWPTLVTTLTYGVQCFARYPDHMKQHDFFKSAMPEG**  **YVQERTISFKDDGTYKTRAEVKFEGDTLVNRIELKGIDFKEDGNILGHKL**  **EYNFNSHNVYITADKQKNGIKANFKIRHNVEDGSVQLADHYQQNTPIGDG**  **PVLLPDNHYLSTQSVLSKDPNEKRDHMVLLEFVTAAGITHGMDELYK*** |
| PhaC-12L-X-G | 12L protease AL-PHA biosensor.  PhaC-fusion amino acid sequence shown. | **MATGKGAAASTQEGKSQPFKVTPGPFDPATWLEWSRQWQGTEGNGHAAAS**  **GIPGLDALAGVKIAPAQLGDIQQRYMKDFSALWQAMAEGKAEATGPLHDR**  **RFAGDAWRTNLPYRFAAAFYLLNARALTELADAVEADAKTRQRIRFAISQ**  **WVDAMSPANFLATNPEAQRLLIESGGESLRAGVRNMMEDLTRGKISQTDE**  **SAFEVGRNVAVTEGAVVFENEYFQLLQYKPLTDKVHARPLLMVPPCINKY**  **YILDLQPESSLVRHVVEQGHTVFLVSWRNPDASMAGSTWDDYIEHAAIRA**  **IEVARDISGQDKINVLGFCVGGTIVSTALAVLAARGEHPAASVTLLTTLL**  **DFADTGILDVFVDEGHVQLREATLGGGAGAPCALLRGLELANTFSFLRPN**  **DLVWNYVVDNYLKGNTPVPFDLLFWNGDATNLPGPWYCWYLRHTYLQNEL**  **KVPGKLTVCGVPVDLASIDVPTYIYGSREDHIVPWTAAYASTALLANKLR**  **FVLGASGHIAGVINPPAKNKRSHWTNDALPESPQQWLAGAIEHHGSWWPD**  **WTAWLAGQAGAKRAAPANYGNARYRAIEPAPGRYVKAKAVLAVIDKRGGG**  **GXXXXXXXGMRKGEELFTGVVPILVELDGDVNGHKFSVRGEGEGDATNGK**  **LTLKFICTTGKLPVPWPTLVTTLTYGVQCFARYPDHMKQHDFFKSAMPEG**  **YVQERTISFKDDGTYKTRAEVKFEGDTLVNRIELKGIDFKEDGNILGHKL**  **EYNFNSHNVYITADKQKNGIKANFKIRHNVEDGSVQLADHYQQNTPIGDG**  **PVLLPDNHYLSTQSVLSKDPNEKRDHMVLLEFVTAAGITHGMDELYK*** |
| PhaC-22L-X-G | 22L protease AL-PHA biosensor.  PhaC-fusion amino acid sequence shown. | **MATGKGAAASTQEGKSQPFKVTPGPFDPATWLEWSRQWQGTEGNGHAAAS**  **GIPGLDALAGVKIAPAQLGDIQQRYMKDFSALWQAMAEGKAEATGPLHDR**  **RFAGDAWRTNLPYRFAAAFYLLNARALTELADAVEADAKTRQRIRFAISQ**  **WVDAMSPANFLATNPEAQRLLIESGGESLRAGVRNMMEDLTRGKISQTDE**  **SAFEVGRNVAVTEGAVVFENEYFQLLQYKPLTDKVHARPLLMVPPCINKY**  **YILDLQPESSLVRHVVEQGHTVFLVSWRNPDASMAGSTWDDYIEHAAIRA**  **IEVARDISGQDKINVLGFCVGGTIVSTALAVLAARGEHPAASVTLLTTLL**  **DFADTGILDVFVDEGHVQLREATLGGGAGAPCALLRGLELANTFSFLRPN**  **DLVWNYVVDNYLKGNTPVPFDLLFWNGDATNLPGPWYCWYLRHTYLQNEL**  **KVPGKLTVCGVPVDLASIDVPTYIYGSREDHIVPWTAAYASTALLANKLR**  **FVLGASGHIAGVINPPAKNKRSHWTNDALPESPQQWLAGAIEHHGSWWPD**  **WTAWLAGQAGAKRAAPANYGNARYRAIEPAPGRYVKAKAGQQSGGSSQGV**  **LAVIDKRGGGGXXXXXXXGMRKGEELFTGVVPILVELDGDVNGHKFSVRG**  **EGEGDATNGKLTLKFICTTGKLPVPWPTLVTTLTYGVQCFARYPDHMKQH**  **DFFKSAMPEGYVQERTISFKDDGTYKTRAEVKFEGDTLVNRIELKGIDFK**  **EDGNILGHKLEYNFNSHNVYITADKQKNGIKANFKIRHNVEDGSVQLADH**  **YQQNTPIGDGPVLLPDNHYLSTQSVLSKDPNEKRDHMVLLEFVTAAGITH**  **GMDELYK*** |
| PhaC-112L-X-G | 112L protease AL-PHA biosensor.  PhaC-fusion amino acid sequence shown. | **MATGKGAAASTQEGKSQPFKVTPGPFDPATWLEWSRQWQGTEGNGHAAAS**  **GIPGLDALAGVKIAPAQLGDIQQRYMKDFSALWQAMAEGKAEATGPLHDR**  **RFAGDAWRTNLPYRFAAAFYLLNARALTELADAVEADAKTRQRIRFAISQ**  **WVDAMSPANFLATNPEAQRLLIESGGESLRAGVRNMMEDLTRGKISQTDE**  **SAFEVGRNVAVTEGAVVFENEYFQLLQYKPLTDKVHARPLLMVPPCINKY**  **YILDLQPESSLVRHVVEQGHTVFLVSWRNPDASMAGSTWDDYIEHAAIRA**  **IEVARDISGQDKINVLGFCVGGTIVSTALAVLAARGEHPAASVTLLTTLL**  **DFADTGILDVFVDEGHVQLREATLGGGAGAPCALLRGLELANTFSFLRPN**  **DLVWNYVVDNYLKGNTPVPFDLLFWNGDATNLPGPWYCWYLRHTYLQNEL**  **KVPGKLTVCGVPVDLASIDVPTYIYGSREDHIVPWTAAYASTALLANKLR**  **FVLGASGHIAGVINPPAKNKRSHWTNDALPESPQQWLAGAIEHHGSWWPD**  **WTAWLAGQAGAKRAAPANYGNARYRAIEPAPGRYVKAKAGQQSGGSSQGG**  **SGSSSGGQGGGSQQSSSGGSQQSQQSQGSGQQSGQQGSQGQQSGGSSQGG**  **SGSSSGGQGGGSQQSSSGGSQQSQQSQGSGQQSGQQGSQVLAVIDKRGGG**  **GXXXXXXXGMRKGEELFTGVVPILVELDGDVNGHKFSVRGEGEGDATNGK**  **LTLKFICTTGKLPVPWPTLVTTLTYGVQCFARYPDHMKQHDFFKSAMPEG**  **YVQERTISFKDDGTYKTRAEVKFEGDTLVNRIELKGIDFKEDGNILGHKL**  **EYNFNSHNVYITADKQKNGIKANFKIRHNVEDGSVQLADHYQQNTPIGDG**  **PVLLPDNHYLSTQSVLSKDPNEKRDHMVLLEFVTAAGITHGMDELYK*** |

**Legend: PhaC**, **Flexible amino acid linker (L)**, **Protease recognition site (X; See Supplementary Table 1), sfGFP (G).**

**Supplementary Table 3. Bacterial strains and constructs.**

| **Strain** | **Relevant features** | **Reference(s)** |
| --- | --- | --- |
| JM109 | endA1, recA1, gyrA96, thi, hsdR17 (r_k_^–^, m_k_^+^), relA1, supE44, Δ(lac-proAB), [F´ traD36, proAB, laqI^q^ZΔM15] | Promega UK |
| EV104-JM109 | JM109 pSB1C3 [EV104]; J23104 promoter and B0034 RBS; BBa_K608002; CamR | This Study  BBa_K608002 |
| C104-JM109 | JM109 pSB1C3-*phaC-phaA-phaB* [C104]; BBa_K1149052; *phaCAB* operon under the control of the J23104 promoter; CamR | This Study  (Kelwick et al., 2015)  BBa_K1149052 |
| pYZW1 | JM109 pSB1C3-*phaC*-12L-G-*phaA-phaB* [PhaC-12L-G]; *phaCAB* operon containing *phaC* fusion gene that includes a 12 amino acid linker and *sfGFP*; CamR | This study |
| pYZW2 | JM109 pSB1C3-*phaC*-12L-T-G-*phaA-phaB* [PhaC-12L-T-G]; *phaCAB* operon containing *phaC* fusion gene that includes a 12 amino acid linker, a TEV protease cleavage site and *sfGFP*; CamR | This study |
| pYZW6 | JM109 pSB1C3-*phaC*-12L-P9-G-*phaA-phaB* [PhaC-12L-P9-G]; *phaCAB* operon containing *phaC* fusion gene that includes a 12 amino acid linker, a MMP9 protease cleavage site and *sfGFP*; CamR | This study |
| pYZW8 | JM109 pSB1C3-*phaC*-12L-P13-G-*phaA-phaB* [PhaC-12L-P13-G]; *phaCAB* operon containing *phaC* fusion gene that includes a 12 amino acid linker, a MMP13 protease cleavage site and *sfGFP*; CamR | This study |
| pYZW9 | JM109 pSB1C3-*phaC*-12L-P14-G-*phaA-phaB* [PhaC-12L-P14-G]; *phaCAB* operon containing *phaC* fusion gene that includes a 12 amino acid linker, a MMP14 protease cleavage site and *sfGFP*; CamR | This study |
| pYZW15 | JM109 pSB1C3-*phaC*-12L-M8-G-*phaA-phaB* [PhaC-12L-M8-G]; *phaCAB* operon containing *phaC* fusion gene that includes a 12 amino acid linker, a ADAM8 protease cleavage site and *sfGFP*; CamR | This study |
| pYZW17 | JM109 pSB1C3-*phaC*-12L-M10-G-*phaA-phaB* [PhaC-12L-M10-G]; *phaCAB* operon containing *phaC* fusion gene that includes a 12 amino acid linker, a ADAM10 protease cleavage site and *sfGFP*; CamR | This study |
| pYZW18 | JM109 pSB1C3-*phaC*-12L-M12-G-*phaA-phaB* [PhaC-12L-M12-G]; *phaCAB* operon containing *phaC* fusion gene that includes a 12 amino acid linker, a ADAM12 protease cleavage site and *sfGFP*; CamR | This study |
| pYZW26 | JM109 pSB1C3-*phaC*-12L-S1-G-*phaA-phaB* [PhaC-12L-S1-G]; *phaCAB* operon containing *phaC* fusion gene that includes a 12 amino acid linker, a ADAMTS1 protease cleavage site and *sfGFP*; CamR | This study |
| pYZW29 | JM109 pSB1C3-*phaC*-12L-S5-G-*phaA-phaB* [PhaC-12L-S5-G]; *phaCAB* operon containing *phaC* fusion gene that includes a 12 amino acid linker, a ADAMTS5 protease cleavage site and *sfGFP*; CamR | This study |
| pYZW33 | JM109 pSB1C3-*phaC*-22L-G-*phaA-phaB* [PhaC-22L-G]; *phaCAB* operon containing *phaC* fusion gene that includes a 22 amino acid linker and *sfGFP*; CamR | This study |
| pYZW34 | JM109 pSB1C3-*phaC*-22L-T-G-*phaA-phaB* [PhaC-22L-T-G]; *phaCAB* operon containing *phaC* fusion gene that includes a 22 amino acid linker, a TEV protease cleavage site and *sfGFP*; CamR | This study |
| pYZW39 | JM109 pSB1C3-*phaC*-112L-G-*phaA-phaB* [PhaC-112L-G]; *phaCAB* operon containing *phaC* fusion gene that includes a 112 amino acid linker and *sfGFP*; CamR | This study |
| pYZW40 | JM109 pSB1C3-*phaC*-112L-T-G-*phaA-phaB* [PhaC-112L-T-G]; *phaCAB* operon containing *phaC* fusion gene that includes a 112 amino acid linker, a TEV protease cleavage site and *sfGFP*; CamR | This study |
| pYZW41 | JM109 pSB1C3-*phaC*-112L-P9-G-*phaA-phaB* [PhaC-112L-P9-G]; *phaCAB* operon containing *phaC* fusion gene that includes a 112 amino acid linker, a MMP9 protease cleavage site and *sfGFP*; CamR | This study |
| pYZW42 | JM109 pSB1C3-*phaC*-112L-P13-G-*phaA-phaB* [PhaC-112L-P13-G]; *phaCAB* operon containing *phaC* fusion gene that includes a 112 amino acid linker, a MMP13 protease cleavage site and *sfGFP*; CamR | This study |
| pYZW43 | JM109 pSB1C3-*phaC*-112L-P14-G-*phaA-phaB* [PhaC-112L-P14-G]; *phaCAB* operon containing *phaC* fusion gene that includes a 112 amino acid linker, a MMP14 cleavage site and *sfGFP*; CamR | This study |
| pYZW44 | JM109 pSB1C3-*phaC*-112L-M8-G-*phaA-phaB* [PhaC-112L-M8-G]; *phaCAB* operon containing *phaC* fusion gene that includes a 112 amino acid linker, a ADAM8 protease cleavage site and *sfGFP*; CamR | This study |
| pYZW45 | JM109 pSB1C3-*phaC*-112L-M10-G-*phaA-phaB* [PhaC-112L-M10-G]; *phaCAB* operon containing *phaC* fusion gene that includes a 112 amino acid linker, a ADAM10 cleavage site and *sfGFP*; CamR | This study |
| pYZW46 | JM109 pSB1C3-*phaC*-112L-M12-G-*phaA-phaB* [PhaC-112L-M12-G]; *phaCAB* operon containing *phaC* fusion gene that includes a 112 amino acid linker, a ADAM12 cleavage site and *sfGFP*; CamR | This study |
| pYZW47 | JM109 pSB1C3-*phaC*-112L-S1-G-*phaA-phaB* [PhaC-112L-S1-G]; *phaCAB* operon containing *phaC* fusion gene that includes a 112 amino acid linker, a ADAMTS1 cleavage site and *sfGFP*; CamR | This study |
| pYZW48 | JM109 pSB1C3-*phaC*-112L-S5-G-*phaA-phaB* [PhaC-112L-S5-G]; *phaCAB* operon containing *phaC* fusion gene that includes a 112 amino acid linker, a ADAMTS5 cleavage site and *sfGFP*; CamR | This study |
| pAJW290 | JM109 pSB1C3-*phaC*-112L-E-G-*phaA-phaB* [PhaC-112L-E-G]; *phaCAB* operon containing *phaC* fusion gene that includes a 112 amino acid linker, a *Schistosoma mansoni* cercarial elastase cleavage site and *sfGFP*; CamR | This study |

**Supplementary Table 4. Oligonucleotide primers.**

| **Number** | **Name** | **Sequence** |
| --- | --- | --- |
| **Primers for cloning** | | |
| RK028 | C104_INF_F | CGCTTGCATGAGTGCCGG |
| RK029 | C104_INF_R | CGCTTTTGCTTTTACATAGCGGC |
| YZW11 | MMP9F | CTGCGCGGCCCGCCGGGCGGCCCGggcatgcgtaaaggcgaagag |
| YZW12 | MMP9R | GCCCGCCGGGCCCGGCGGGCCCGGaccgccaccgccacg |
| YZW19 | MMP13F | GCGGATGCGGATggcatgcgtaaaggcgaagag |
| YZW20 | MMP13R | CACCAGGCCCGCCGGaccgccaccgccacg |
| YZW21 | MMP14F | GGCCCGCCGAACGGCATGCGTAAAGGCGAAGAG |
| YZW22 | MMP14R | CACCAGGCCCGCCGGACCGCCACCGCCACG |
| YZW37 | ADAM8F | CAGCGCCTGCGCggcatgcgtaaaggcgaagag |
| YZW38 | ADAM8R | CGCCGCCGCCGGaccgccaccgccacg |
| YZW41 | ADAM10F | GTGGTGAGCCAGGGCATGCGTAAAGGCGAAGAG |
| YZW42 | ADAM10R | CGCCTGCGCGCTACCGCCACCGCCACG |
| YZW43 | ADAM12F | CGCACCCCGAGCGATAAAggcatgcgtaaaggcgaagag |
| YZW44 | ADAM12R | GCTGCTGCTGCGCACCGCaccgccaccgccacg |
| YZW59 | ADAMTS1F | GCGCGCGGCGGCggcatgcgtaaaggcgaagag |
| YZW60 | ADAMTS1R | TTCCTGCGCCACaccgccaccgccacg |
| YZW65 | ADAMTS5F | CCGCGGCGGCATGGTGggcatgcgtaaaggcgaagag |
| YZW66 | ADAMTS5R | CCTTCCGGCGCCGGaccgccaccgccacg |
| YZW77 | TEV-F | TATTTTCAGGGCGGCATGCGTAAAGGCGAAGAG |
| YZW78 | TEV-R | CAGGTTTTCACCGCCACCGCCACG |
| YZW81 | 22aa-F | GGCTCAAGTCAGGGGGTGCTGGCGGTGATTGATAAACGTGGC |
| YZW82 | 22aa-R | GCCTGACTGCTGCCCCGCTTTTGCTTTTACATAGCGGCC |
| YZW83 | BIO-INF-F | GTGCTGGCGGTGATTGATAAACGT |
| YZW84 | BIO-INF-R | CGCTTTTGCTTTTACATAGCGGCC |
| AJW671 | 5-phaC-ELA | cCActtggcatgcgtaaaggcgaagag |
| AJW672 | 3-phaC-ELA | CCAACTACCGCCACCGCCACGTTTATC |
| **Primers for sequencing** | | |
| RK013 | VF2 | TGCCACCTGACGTCTAAGAA |
| RK014 | VR | ATTACCGCCTTTGAGTGAGC |
| RK015 | PhaCseq1 | TGGCAGGCGATGGCGGAAG |
| RK016 | PhaB_F | ATGACTCAGCGCATTGCGTATGTGA |
| RK017 | PhaCseq2 | CTATGGCAACGCGCGCTACC |
| RK019 | PhaAseq1 | GTTCCCTCCCGTTTCC |
| RK020 | PhaBseq1 | GCGACGATAACGAAGCC |
| RK023 | PhaC_seq4_F | TCTGCTGCGCGGTCTGG |
| RK026 | PhaA2_FWD | GTTGACGGCTTATGGGATGT |
| RK027 | GFP_F | AAGAGCTGTTCACTGGTGTCG |

**Supplementary Table 5. High-throughput metalloproteinase AL-PHA biosensor screening assay data**

| **AL-PHA Biosensor** | **Supernatant**  **(relative fold change / %)** | **Supernatant**  **t-test p-value** | **AL-PHA bead**  **(relative fold change / %)** | **AL-PHA bead**  **t-test p-value** |
| --- | --- | --- | --- | --- |
| Control (PhaC-112L-G) | 0.25±0.44-fold / NC | n.s. p=0.60 | +0.08±0.13-fold / NC | n.s. p=0.60 |
| TEV (PhaC-112L-T-G) | 0.11±0.36-fold / NC | n.s. p=0.77 | -0.03±0.11-fold / NC | n.s. p=0.81 |
| MMP9 (PhaC-112L-P9-G) | 0.22±0.09-fold / ~22% increase | n.s. p=0.07 | -0.27±0.18-fold / ~27% decrease | n.s. p=0.21 |
| MMP13 (PhaC-112L-P13-G) | 1.13±0.30-fold / ~113% increase | *p=0.02 | -0.22±0.05-fold / ~22% decrease | *p=0.01 |
| MMP14 (PhaC-112L-P14-G) | 0.81±0.25-fold / ~81% increase | *p=0.03 | -0.01±0.06-fold / NC | n.s. p=0.89 |
| ADAM8 (PhaC-112L-M8-G) | 1.14±0.25-fold / ~114% increase | *p=0.01 | -0.17±0.07-fold / ~17% decrease | n.s. p=0.08 |
| ADAM10 (PhaC-112L-M10-G) | 0.99±0.22-fold / ~99% increase | *p=0.01 | -0.45±0.21-fold / ~45% decrease | n.s. p=0.09 |
| ADAM12 (PhaC-112L-M12-G) | 0.80±0.25-fold / ~80% increase | *p=0.03 | -0.33±0.12-fold / ~33% decrease | p=0.05 |
| ADAMTS-1 (PhaC-112L-S1-G) | 1.01±0.47-fold / ~101% increase | n.s. p=0.097 | -0.22±0.10-fold / ~22% decrease | n.s. p=0.11 |
| ADAMTS-5 (PhaC-112L-S5-G) | 1.15±0.33-fold / ~115% increase | *p-0.03 | -0.03±0.30-fold / NC | n.s. p=0.93 |

Data is shown in Figure 6 of the main manuscript, Student *t*-test not significant (n.s.) or *P<0.05. NC = Due to the relatively large error between replicates there is no clear (NC) increase in supernatant fluorescence or decrease in AL-PHA bead fluorescence.

**Supplementary Table 6. Comparative advantages and disadvantages of AL-PHA beads**

|  | **AL-PHA beads** | **Small molecule substrates** | **Peptides** | **Quantum dots** | **FRET-probes** | **Whole-cell bioreporters** |
| --- | --- | --- | --- | --- | --- | --- |
| **Modular Design** | √√√ | √ | √√ | √ | √√√ | √√√ |
| **Development time-scale** | Several days | Several weeks | Several days or weeks | Several weeks | Several days | Several days |
| **Assay flexibility** | √√√ | √√√ | √√√ | √√ | √√√ | √ |
| **Producible within a standard molecular laboratory** | YES | POSSIBLY | POSSIBLY | NO | YES | YES |
| **Shelf-life** | √√√ | √√√ | √√ | √ | √√ | √√ |
| **Regulated GMO** | No | No | No | No | No | Potentially |
| **Production cost** | £ | £ | ££ | £££ | ££ | £ |
| **Environmentally friendly** | √√√ | √ | √√√ | √ | √√√ | √√ |

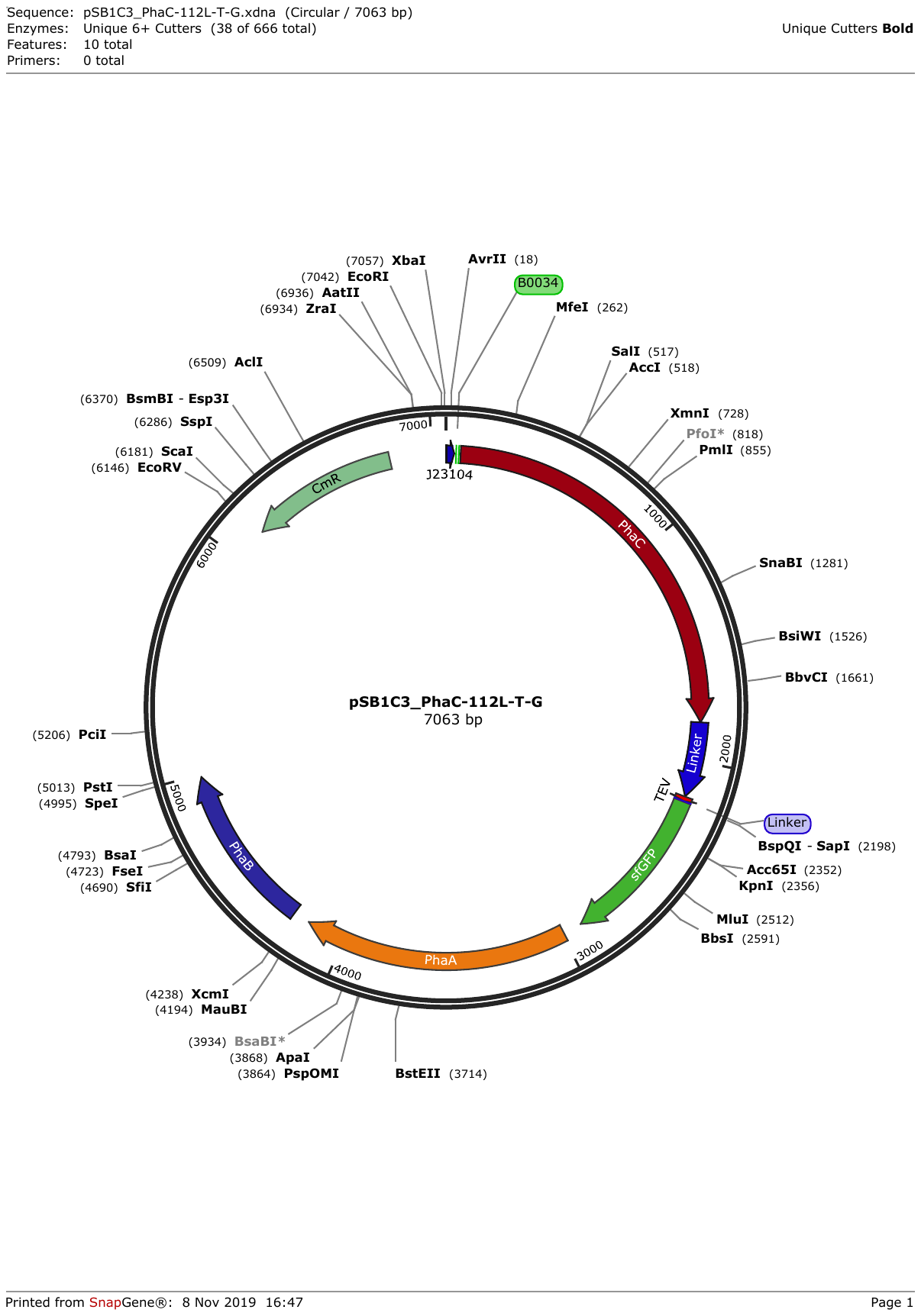

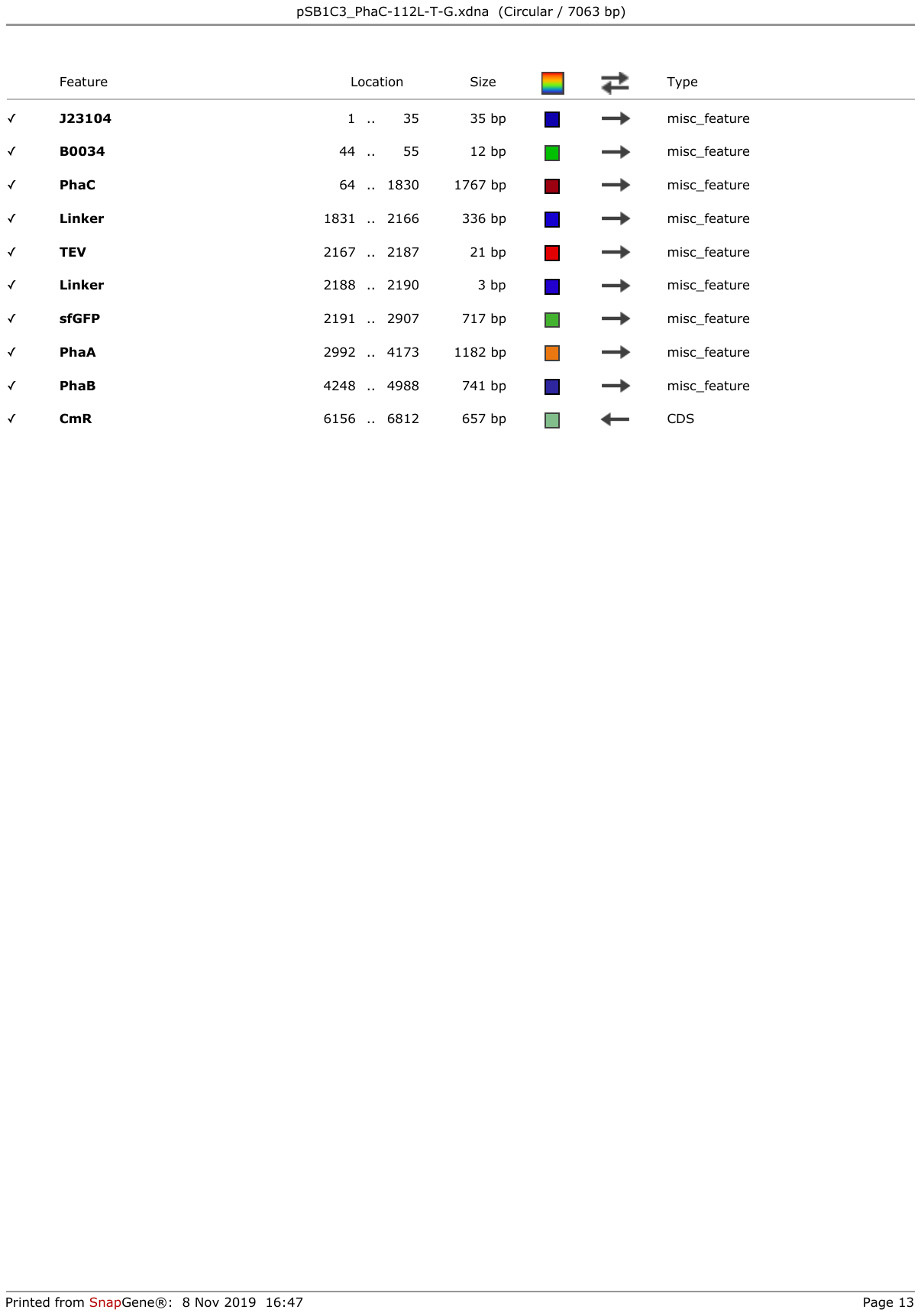

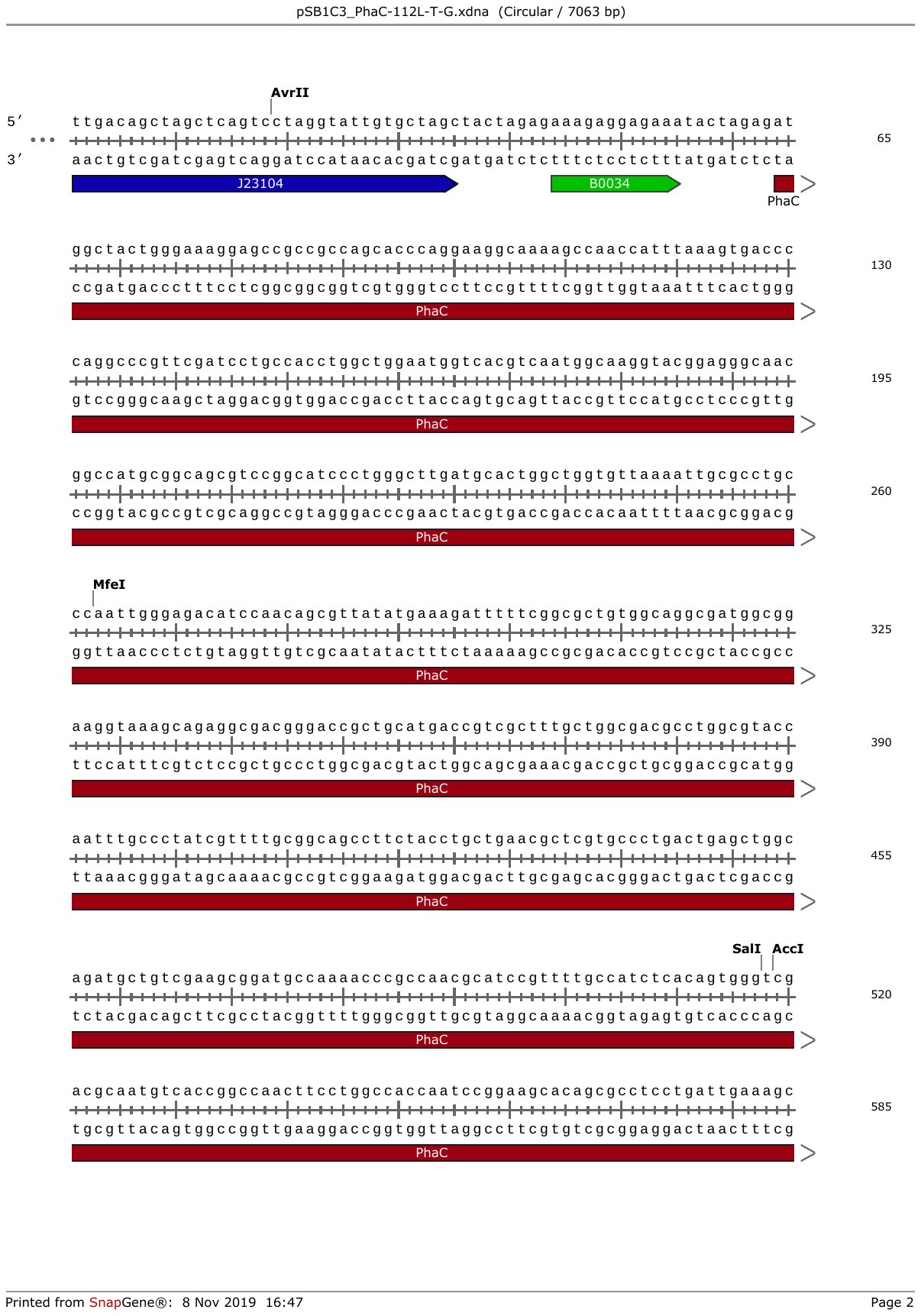

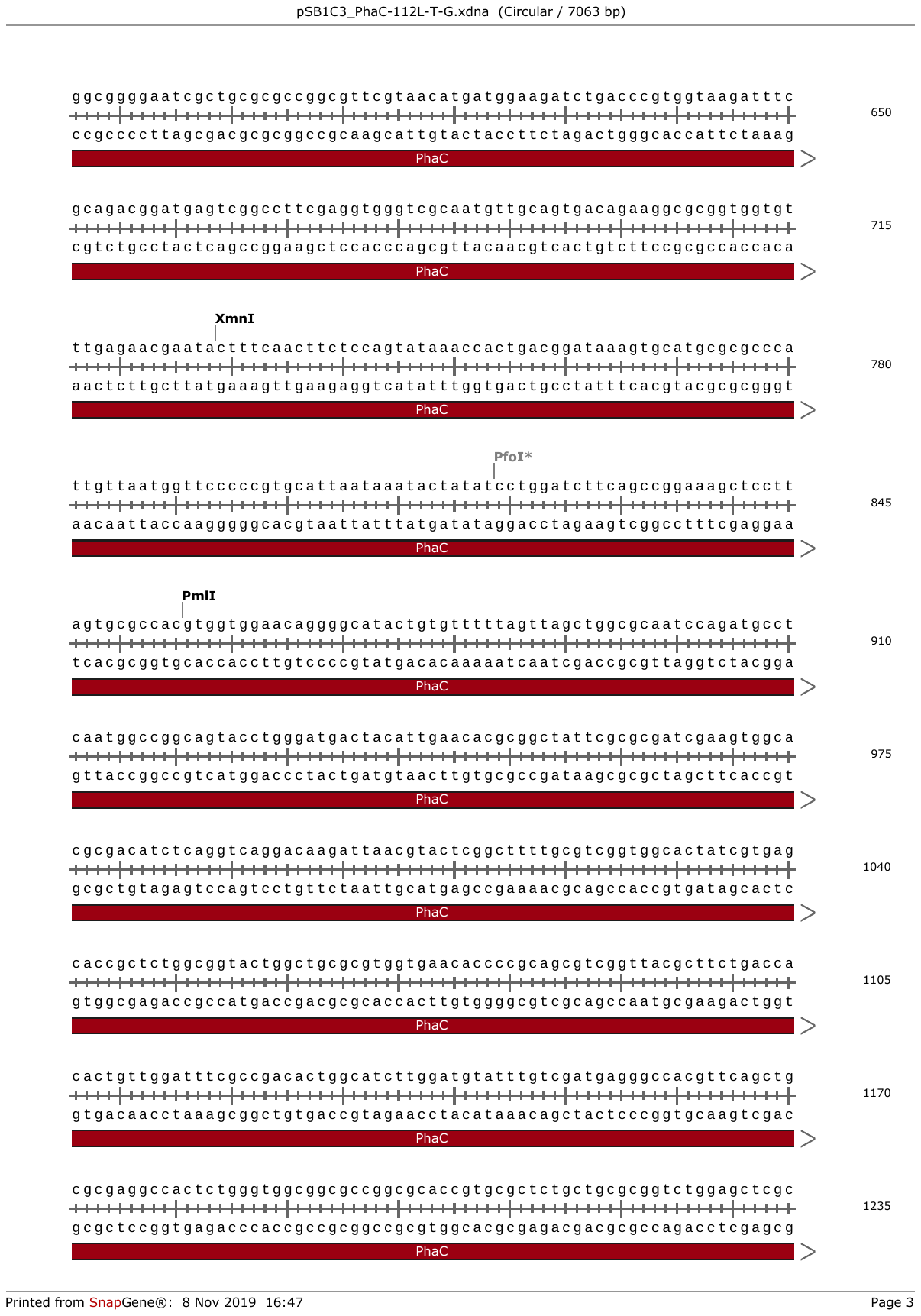

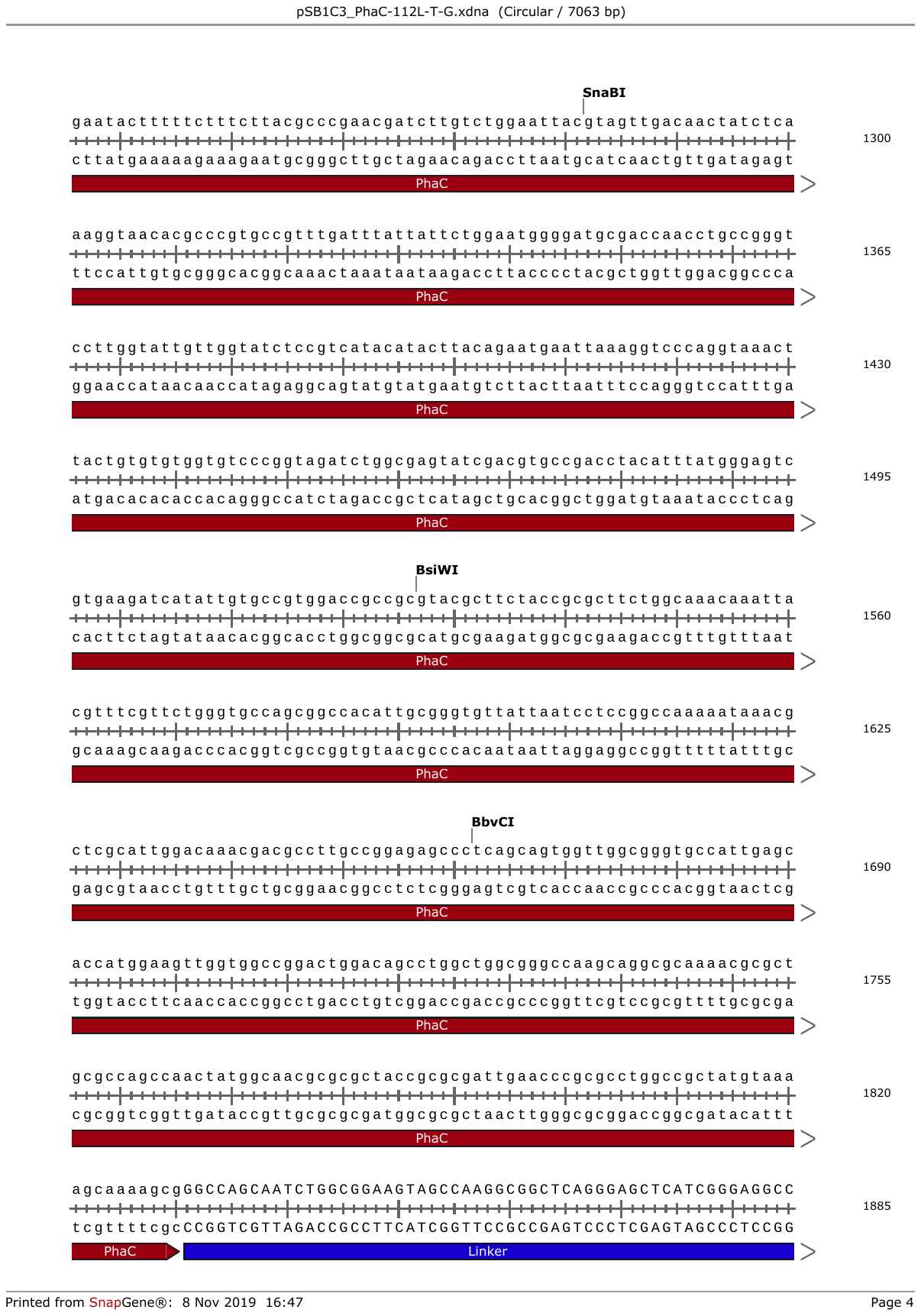

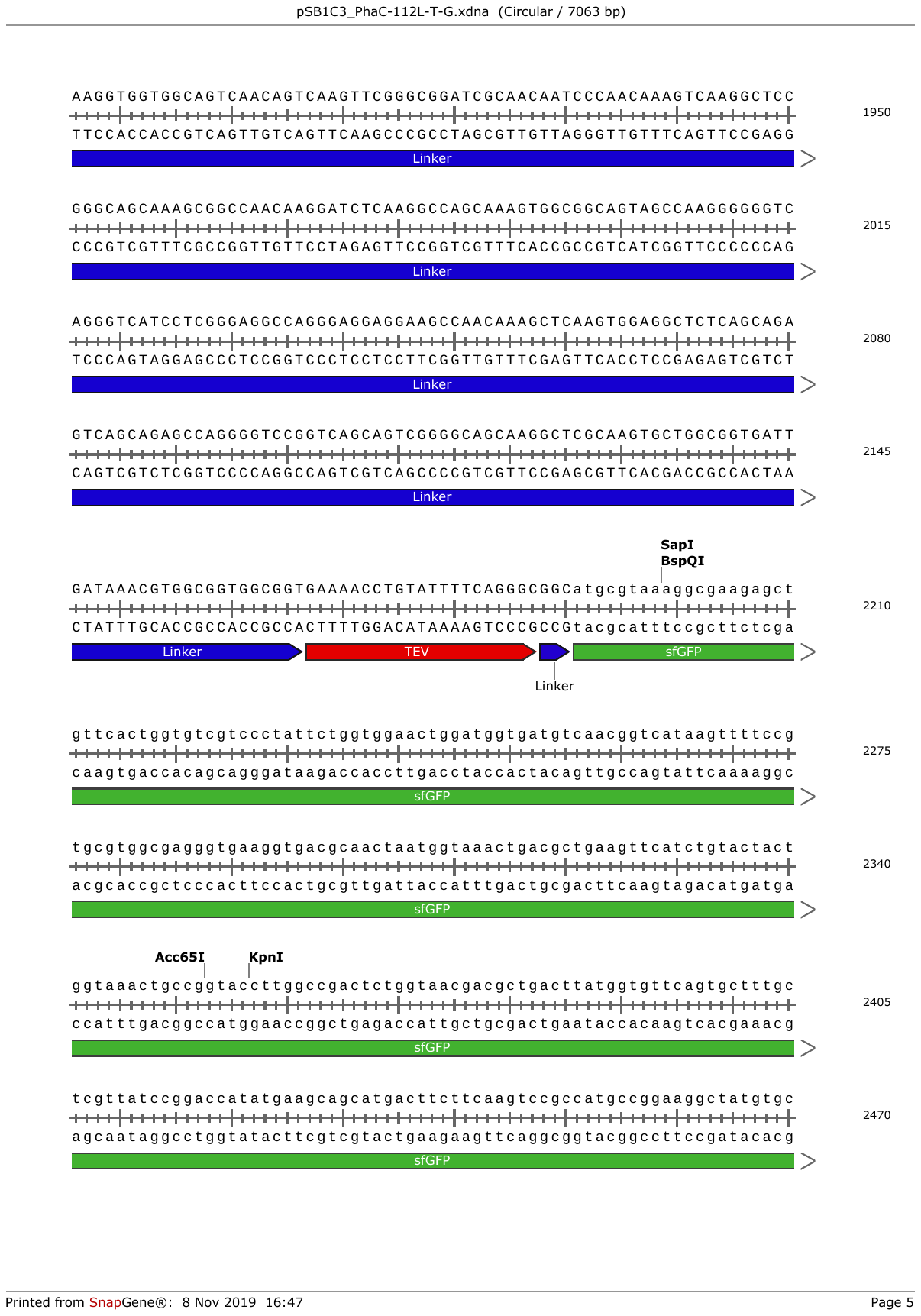

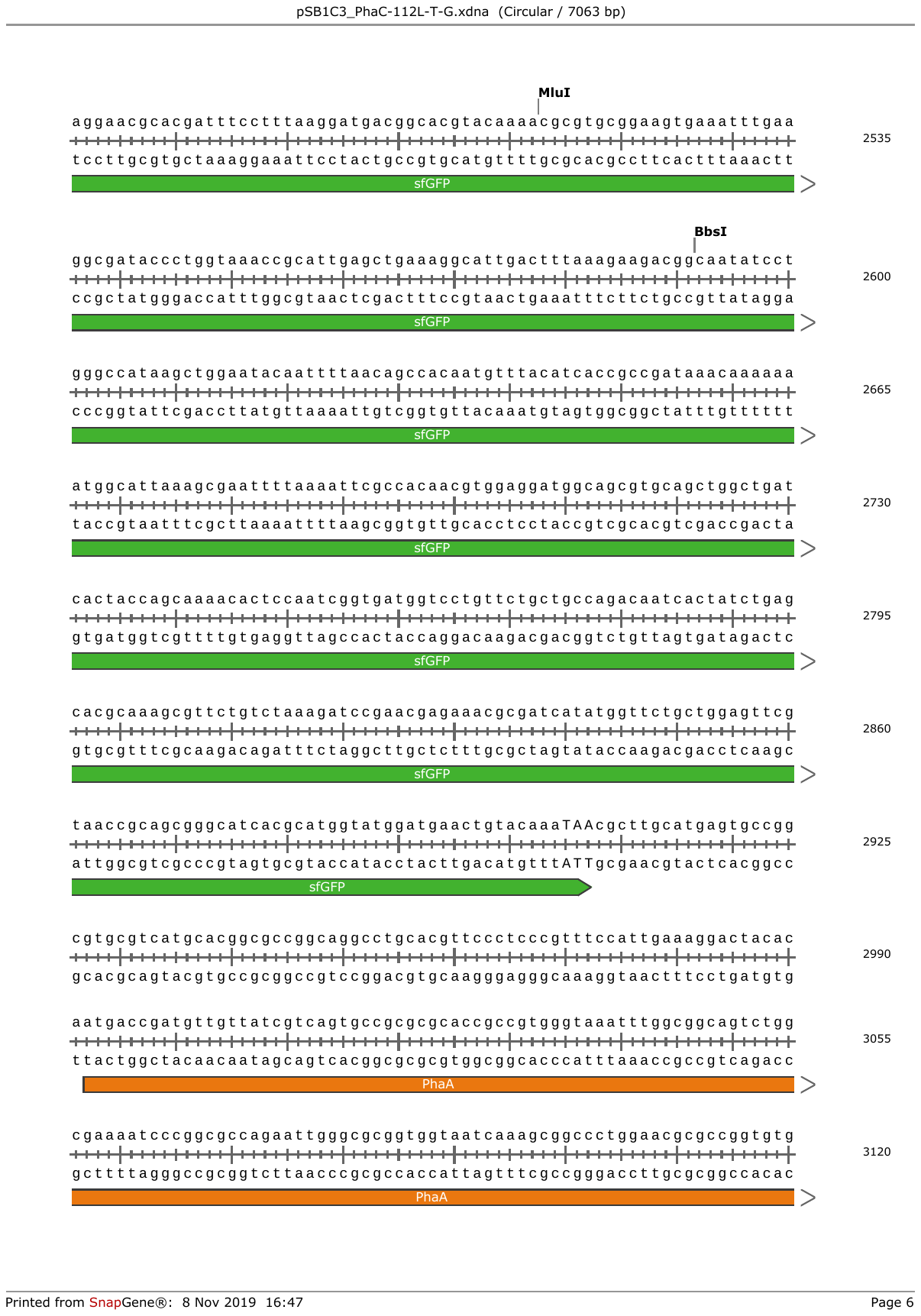

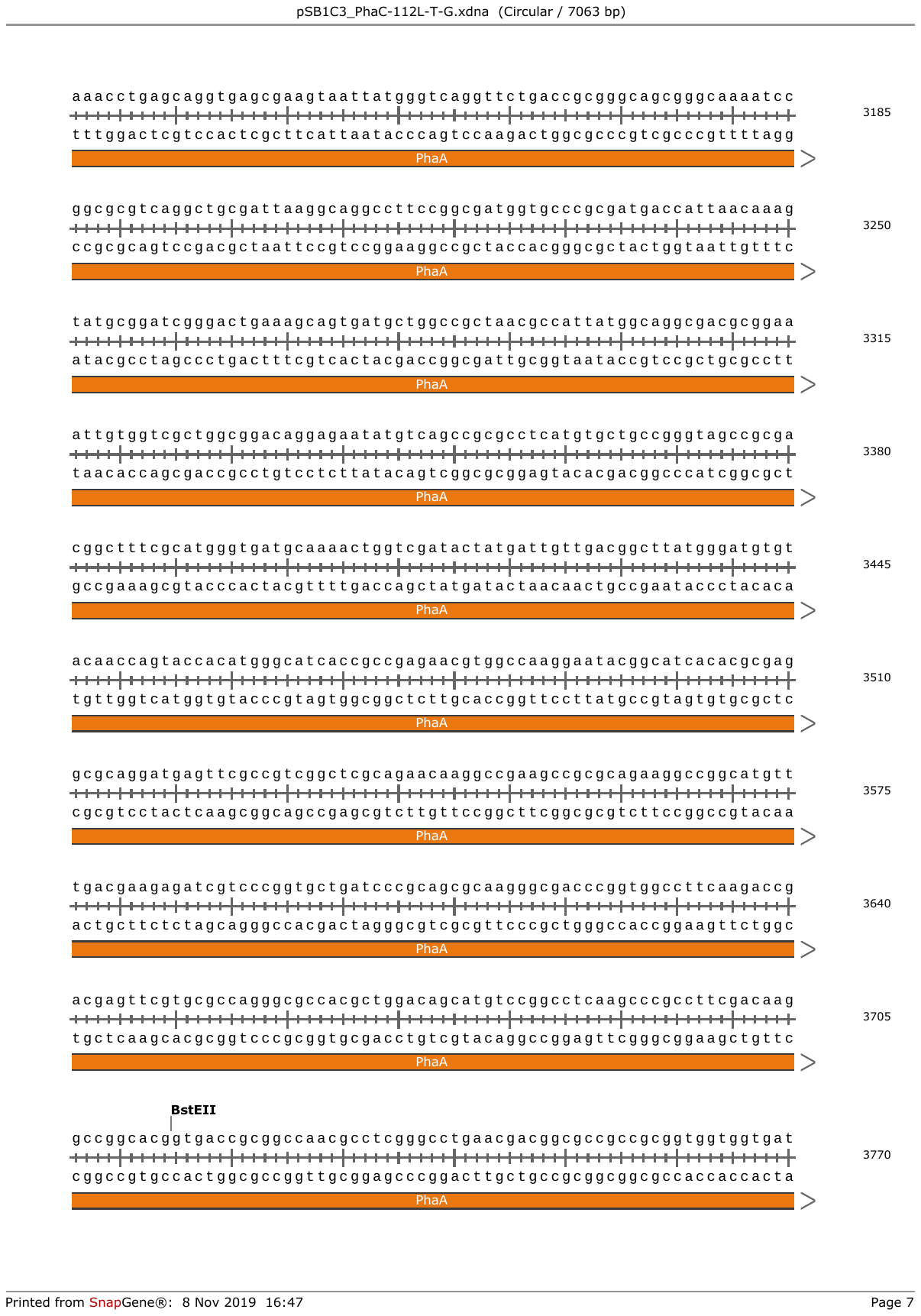

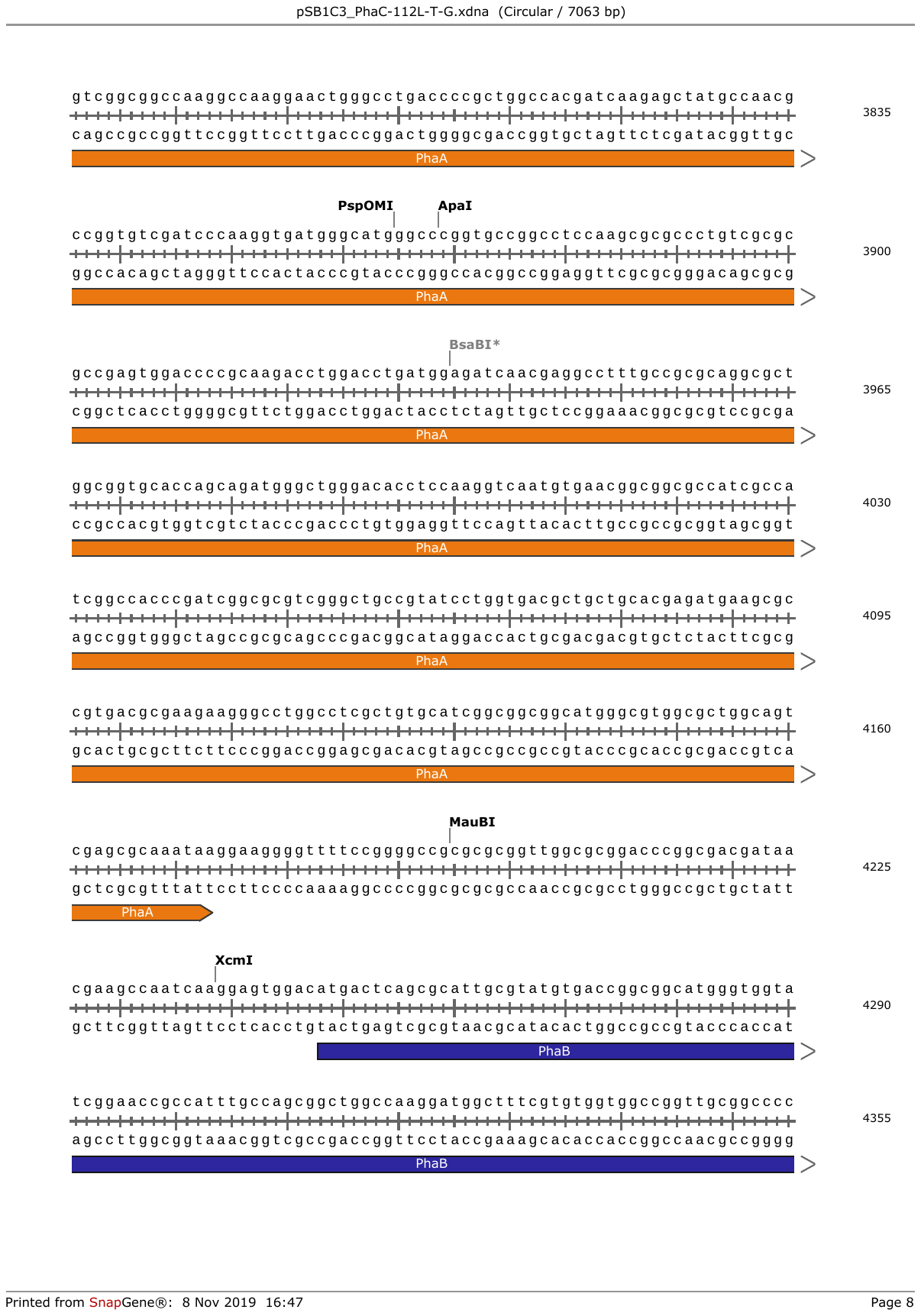

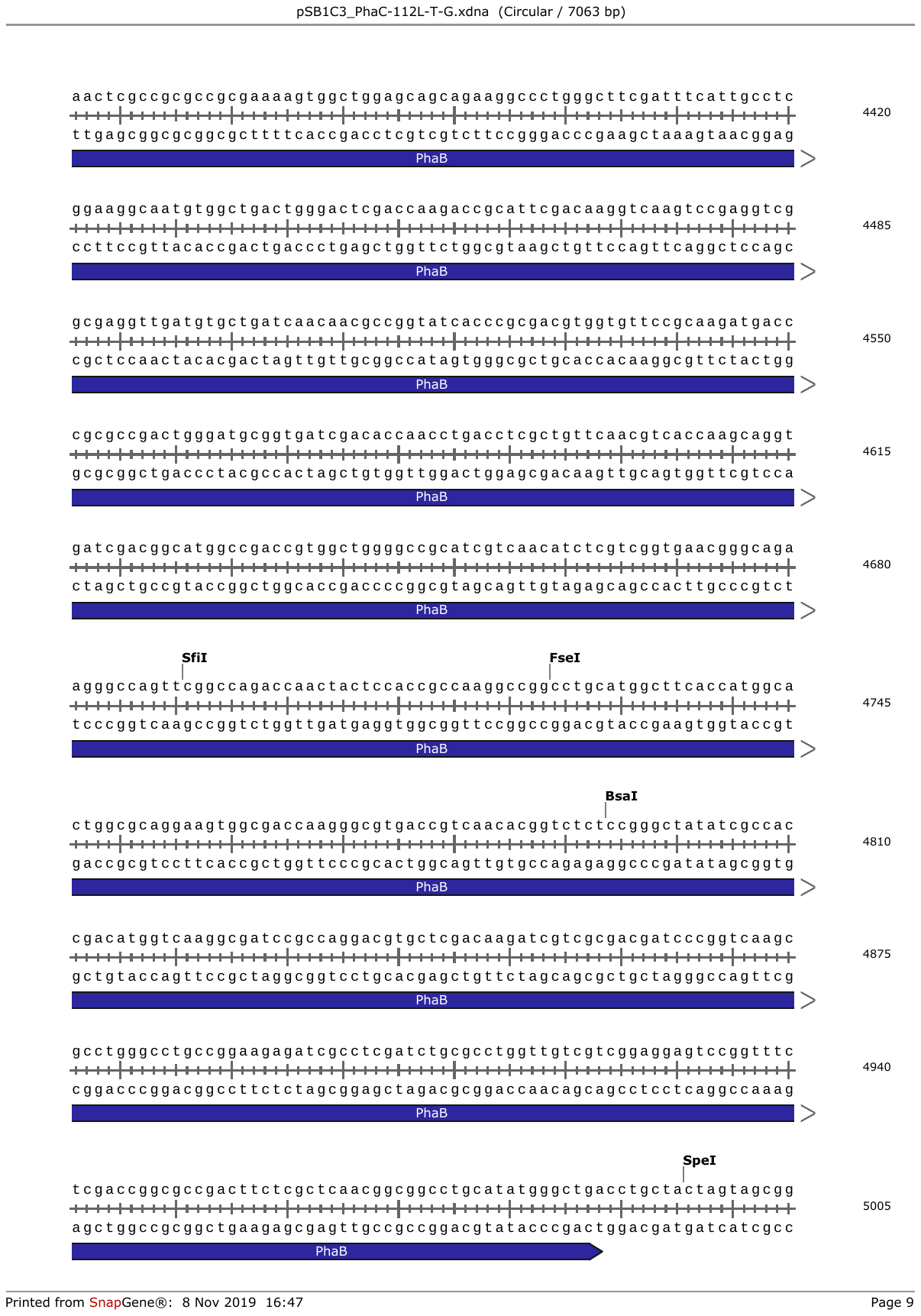

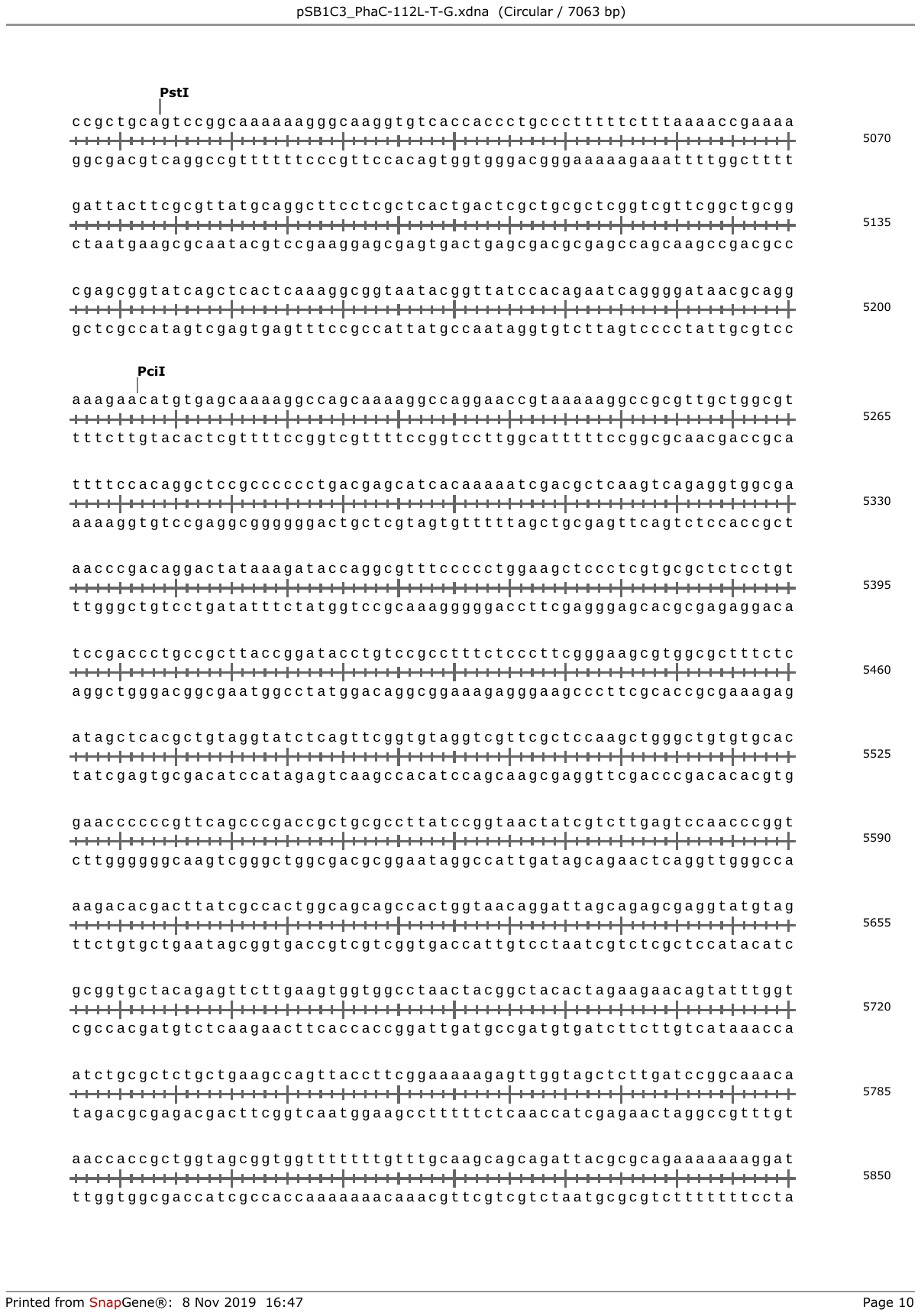

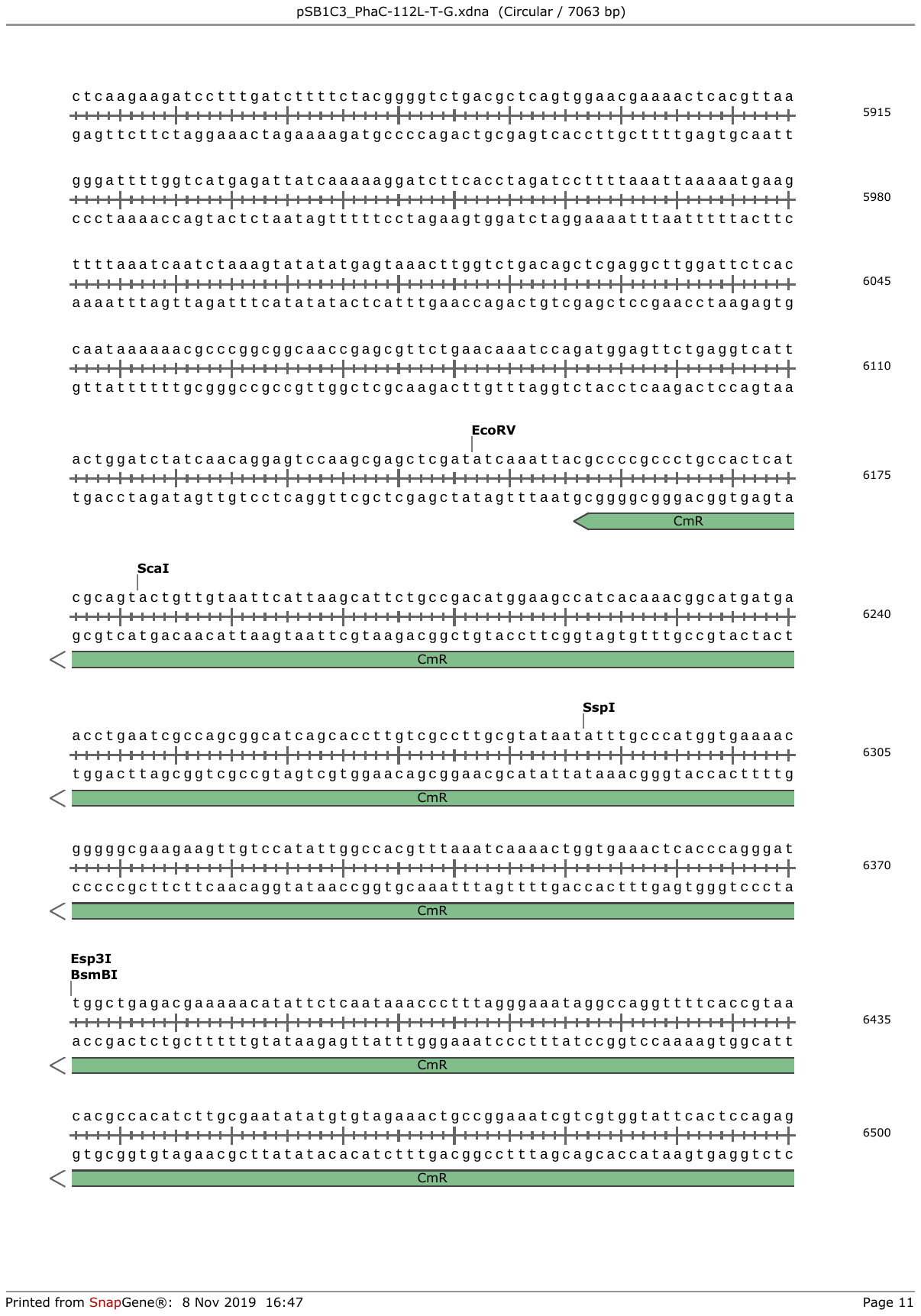

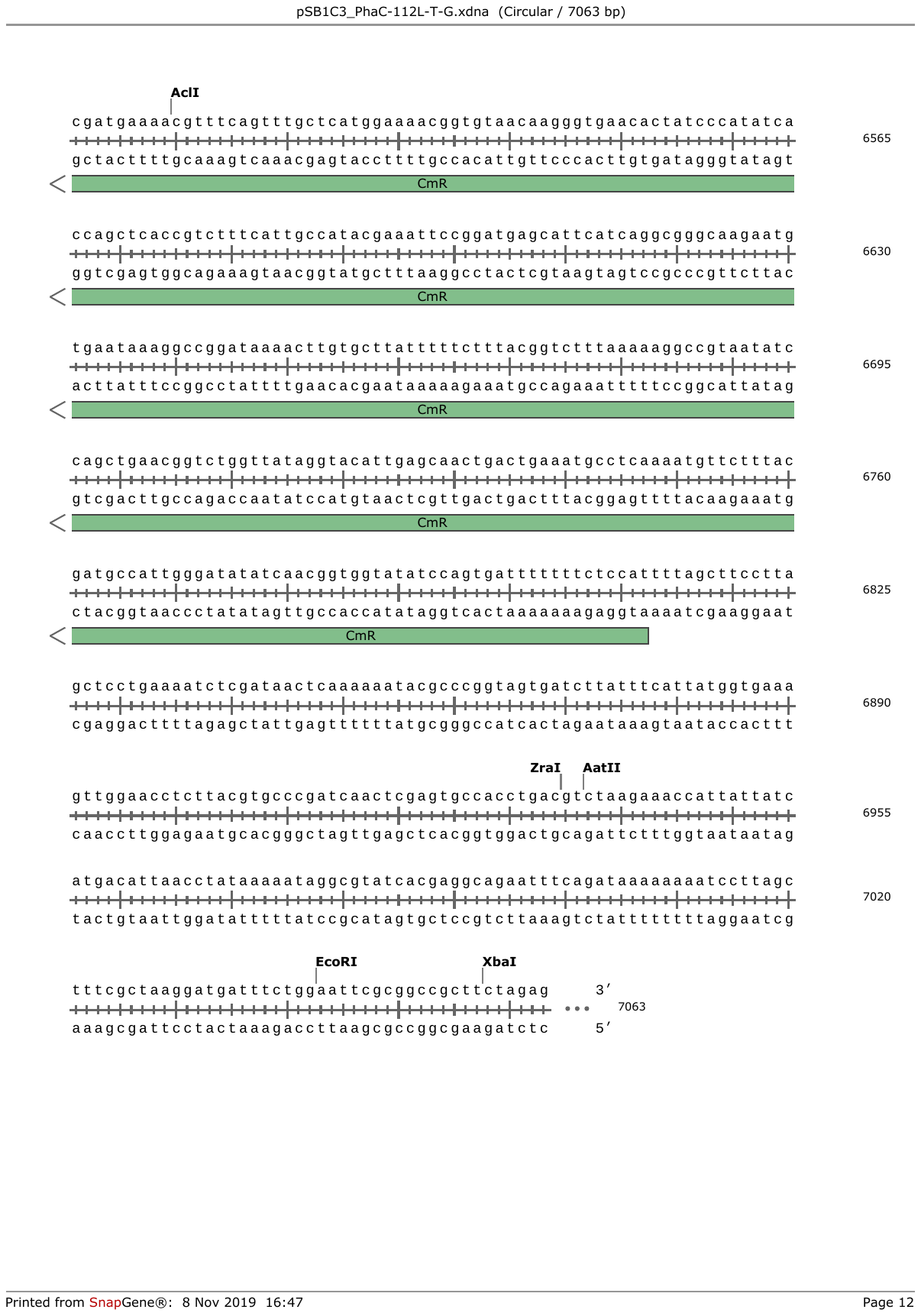

**Supplementary Figure 1. Third generation AL-PHA TEV biosensor plasmid map and sequence.** Plasmid map and sequence of plasmid pYZW40 for production of PhaC-112L-T-G AL-PHA biosensor beads. Plasmid map generated using SnapGene Viewer software v5.1.1.

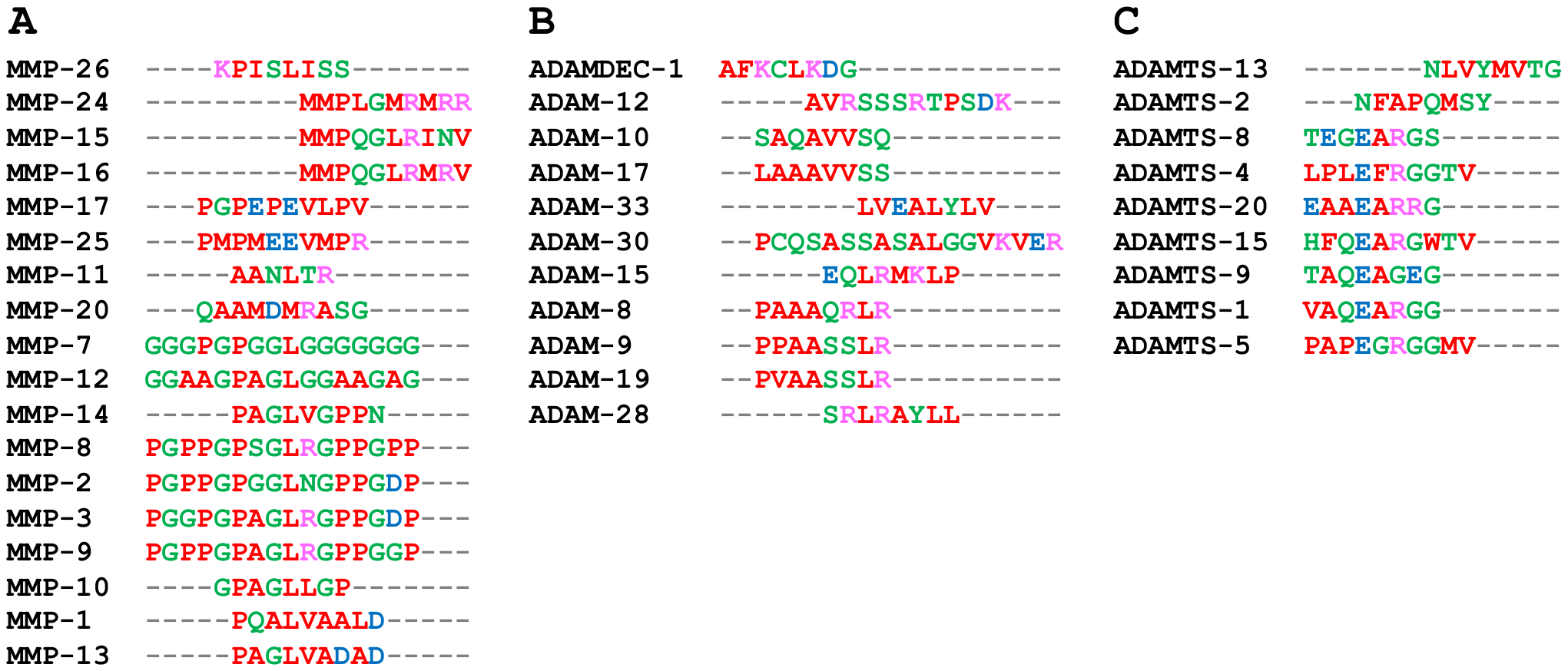

**Supplementary Figure 2. Multiple sequence alignments of metalloproteinase recognition sites.** Protease recognition sites for **(A)** Matrix metalloproteinases [MMPs], **(B)** A disintegrin and metalloproteinases [ADAMs] and **(C)** A disintegrin and metalloproteinase with thrombospondin motifs [ADAMTSs] were identified from the literature (see Supplementary Table 1) and were aligned using Clustal Omega v1.2.4 (<https://www.ebi.ac.uk/Tools/msa/clustalo/>).

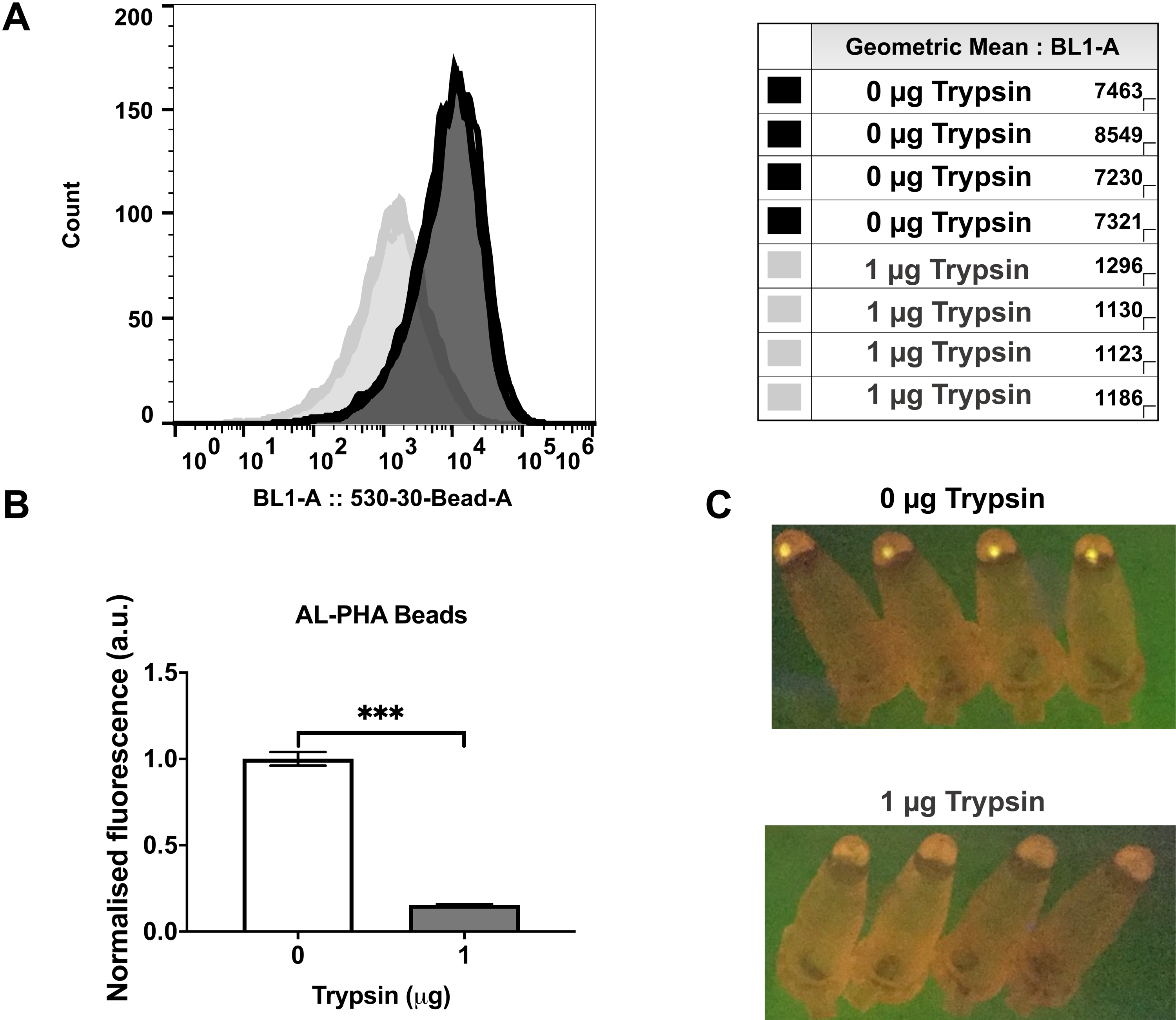

**Supplementary Figure 3. Trypsin digest of AL-PHA biosensors.** AL-PHA biosensor beads (PhaC-22L-G) were treated with either 0 μg or 1 μg of Trypsin for 2h, at 37^o^C with 220 rpm shaking (Eppendorf, Thermo Mixer C). Post trypsin assay, AL-PHA biosensor bead samples were analysed using flow cytometry (Attune NxT). **(A)** Flow cytometry fluorescence histogram showing all experimental replicates and their sample fluorescence data. **(B)** Geometric means (BL1-A, Ex 488 nm Em. 530-30 nm) of trypsin treated AL-PHA biosensor bead samples were normalised against untreated samples of the same bead batch. Error bars indicate the standard error of the mean of four independent replicates (different AL-PHA biosensor bead batches). Student t-test, ***P**<**0.001. **(C)** Post trypsin assay, samples were centrifuged (6000 *g*) and pelleted AL-PHA biosensor bead samples were imaged on a Visi-Blue transilluminator (Analytik Jena, USA) using an iPhone 6 (Apple Inc., USA) camera.

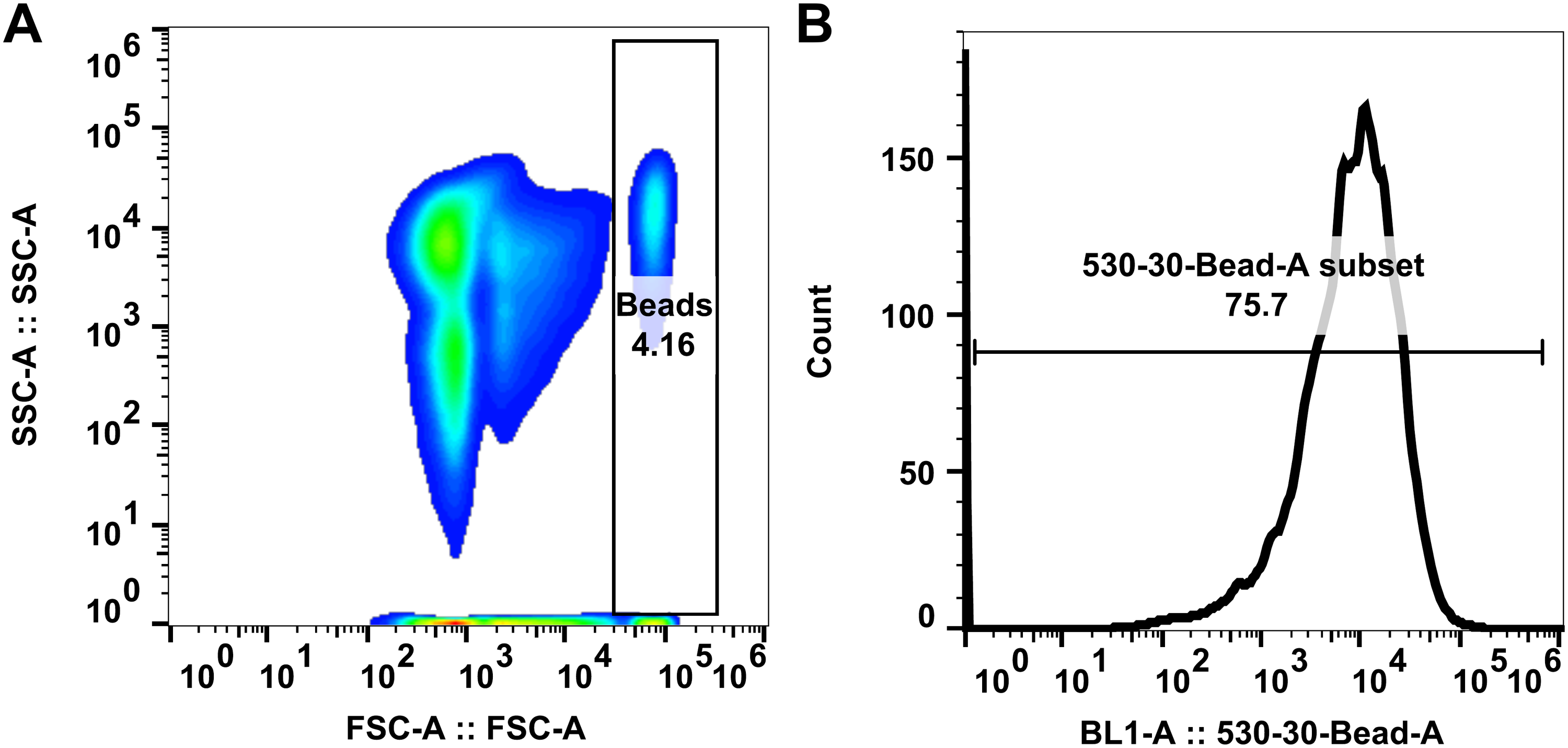

**Supplementary Figure 4. Flow cytometry gating strategies used for AL-PHA biosensor bead analyses. (A)** Representative forward scatter (FSC-A) and side scatter (SSC-A) contour plot, including the flow cytometry gating strategy (Beads) used for AL-PHA biosensor bead analyses. **(B)** Representative fluorescence histogram of AL-PHA biosensor beads (PhaC-22L-G), including the additional flow cytometry gating strategy (Bead-A subset) that was used to remove flow cytometry equipment signal noise (Attune NxT; BL1-A, Ex 488 nm Em. 530-30 nm) from AL-PHA biosensor bead analyses.

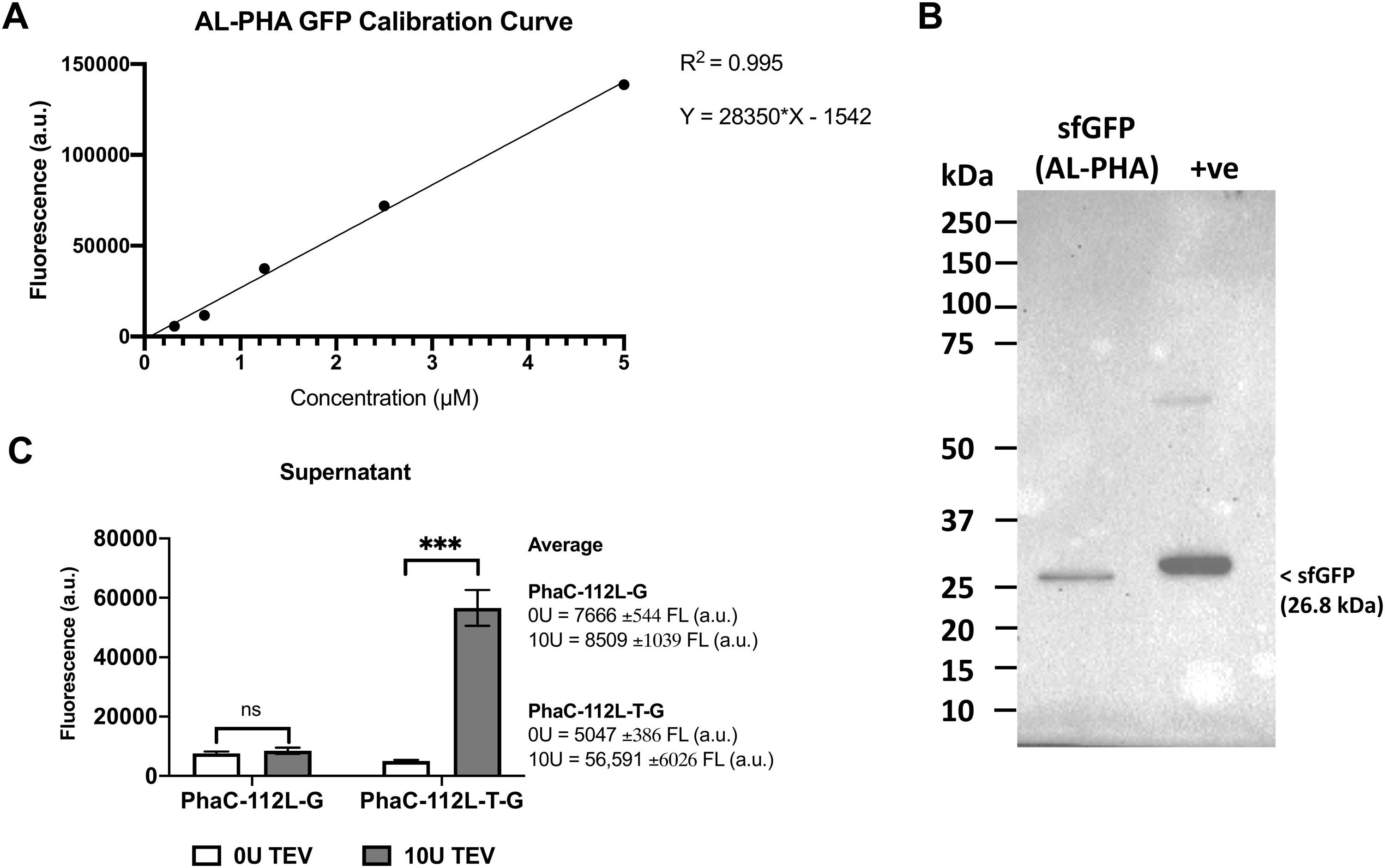

**Supplementary Figure 5. GFP Calibration Curve. (A)** In order to convert sfGFP fluorescence (a.u.) in the assay supernatant into protein concentration (µM) a sfGFP calibration curve was setup using a range of purified sfGFP protein concentrations. The sfGFP used to create this calibration curve was proteolytically released (purified) from several batches of third generation TEV AL-PHA biosensor (PhaC-112L-T-G) reactions (n=4). **(B)** Western blot analysis (anti-GFP antibody, 1:4,000 dilution, #A10260, Thermo Fisher Scientific Ltd, UK) of sfGFP protein used in the GFP calibration curve. Recombinant GFPmut3b was used as a positive control **(C)** AL-PHA assay data from Figure 1F in the main text. Raw fluorescence values are shown (483-14 nm/530-30nm, 2000 gain, BMG ClARIOstar plate reader) to enable calculation of sfGFP released into the assay supernatant during the AL-PHA assay (PhaC-112L-T-G, 10 U AcTEV; 1.87 μM sfGFP equals ~10-fold increase in supernatant fluorescence). Error bars denote standard error of the mean, n=4, Student *t*-test ***P<0.001.

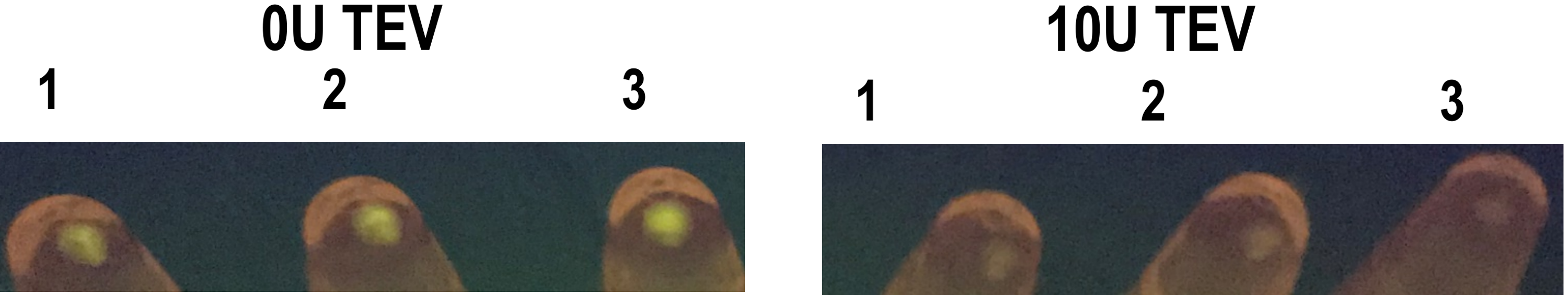

**Supplementary Figure 6. AcTEV treatment removes sfGFP. AL-PHA beads.** AL-PHA biosensor beads (PhaC-112L-T-G) were treated with either 0 U or 10 U of Tobacco Etch Virus protease (AcTEV, ThermoFisher #12575015) for 2h, at 30^o^C with 500 rpm shaking (Eppendorf, Thermo Mixer C). Post assay, samples were centrifuged (6000 *g*) and pelleted AL-PHA biosensor bead samples were imaged on a Visi-Blue transilluminator (Analytik Jena, USA) using an iPhone 6 (Apple Inc., USA) camera.

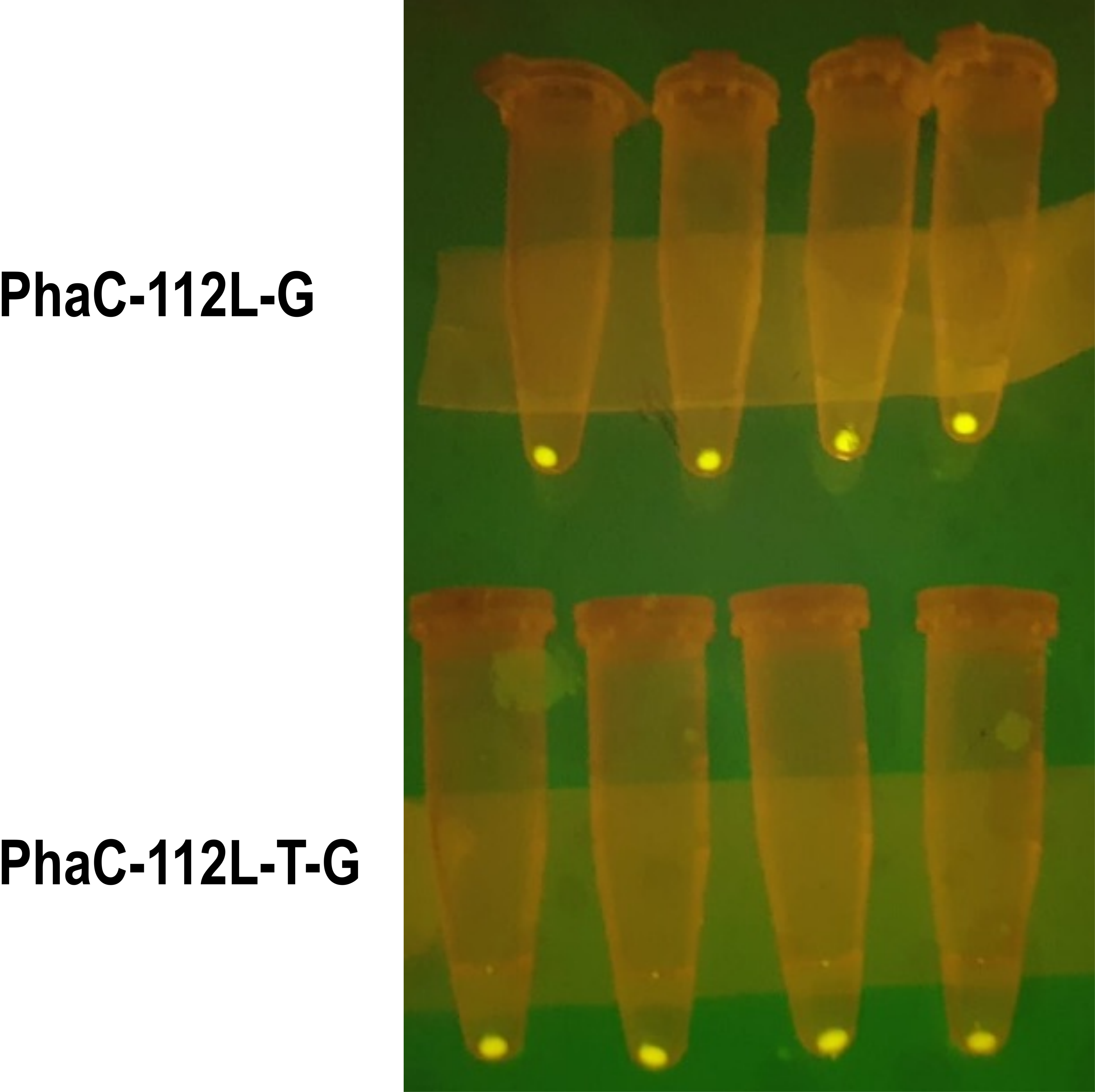

**Supplementary Figure 7. AL-PHA biosensor beads are fluorescent 12-months post-production.** Twelve-month-old AL-PHA biosensor bead batches (PhaC-112L-G and PhaC-112L-T-G), stored at 4^o^C, were diluted (1:50) into PBS (1X) and centrifuged (6000 *g*). Pelleted AL-PHA biosensor bead samples were imaged on a Visi-Blue transilluminator (Analytik Jena, USA) using an iPhone 6 (Apple Inc., USA) camera.

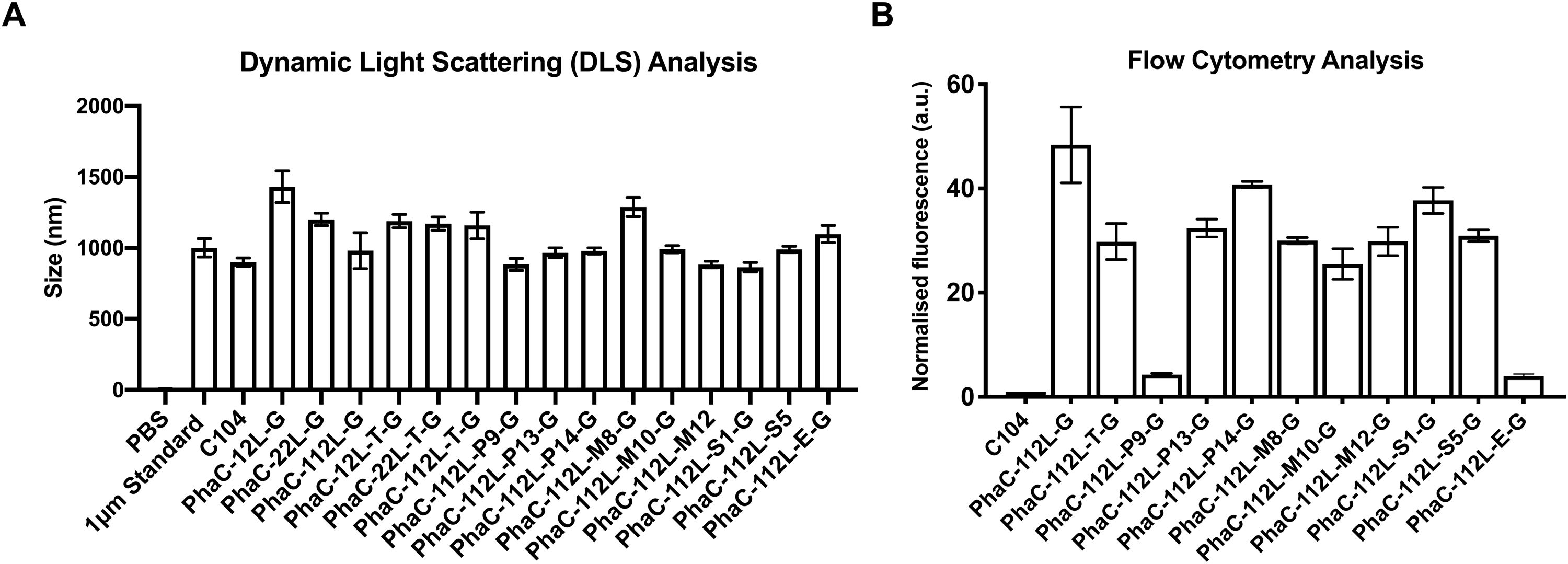

**Supplementary Figure 8. Characterisation of AL-PHA biosensor beads. (A)** Bead sizes were determined using dynamic light scattering (DLS). PBS (alone), commercial 1 μm standard beads (ThermoFisher, USA #F13839), non-functionalised control PHA beads (C104) and AL-PHA biosensor beads were diluted (1:100) into 1X PBS and analysed using a Zetasizer Nano ZS (Malvern Instruments, UK). **(B)** Surface functionalisation (sfGFP fluorescence) of AL-PHA biosensor beads was analysed using flow cytometry (Attune NxT). Geometric means (BL1-A, Excitation 488 nm Emission 530-30 nm) of AL-PHA biosensor bead samples were normalised against the average geometric mean of C104 control bead batches. Error bars indicate the standard error of the mean of 3-4 independent replicates (different C104 or AL-PHA biosensor bead batches).

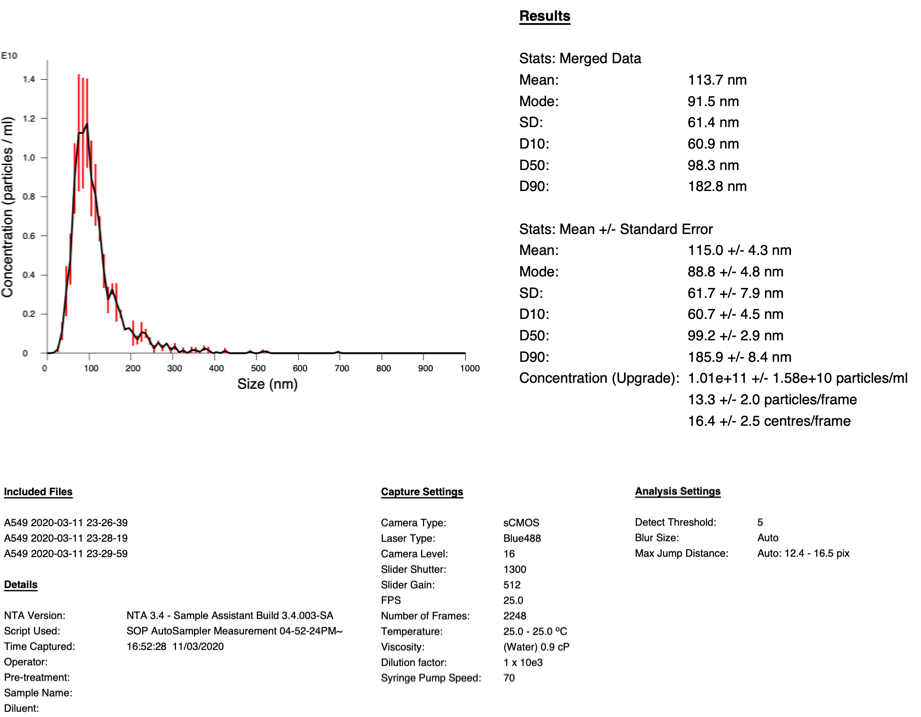

**Supplementary Figure 9. Nanoparticle tracking analysis (NTA) of A549 extracellular vesicles.** A549 cell line EVs (#HBM-A549-100; HansaBioMed, Estonia) were diluted 1:1000 into PBS (1X) and analysed using an NS300 NTA system equipped with a 96-well plate autosampler (Malvern Instruments, UK).

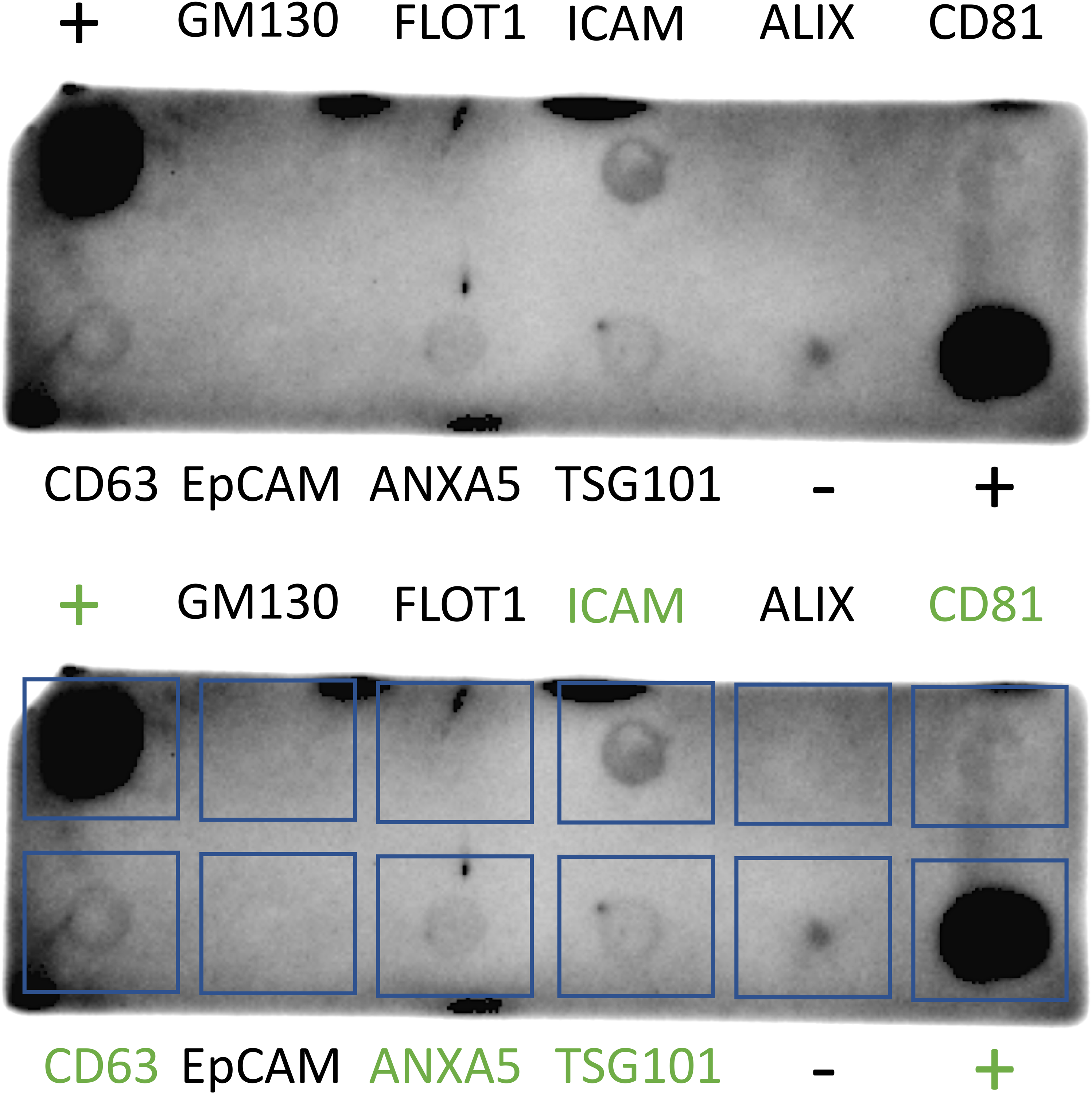

**Supplementary Figure 10. Exo-Check Exosome Antibody Array analysis of A549 extracellular vesicles.** 100 μg of A549 cell line EVs (#HBM-A549-100; HansaBioMed, Estonia) were processed, according to manufacturer’s instructions, and analysed for the presence of EV markers using the Exo-Check Exosome Antibody Array (System Biosciences, LLC., USA). The Exo-Check array included 12 dot-blot spots and was used with antibodies specific for the exosome markers CD63, CD81, ALIX, FLOT1, ICAM1, EpCam, ANXA5 and TSG101, as well as several cellular contamination controls (GM130 cis-Golgi marker) and several assay positive/negative controls. The developed dot blot array was imaged using a ChemiDoc imaging system (Bio-Rad Laboratories Inc., USA). Top and bottom panels are the same dot blot array. Blue boxes added to highlight array dot areas and green text indicates positive results.

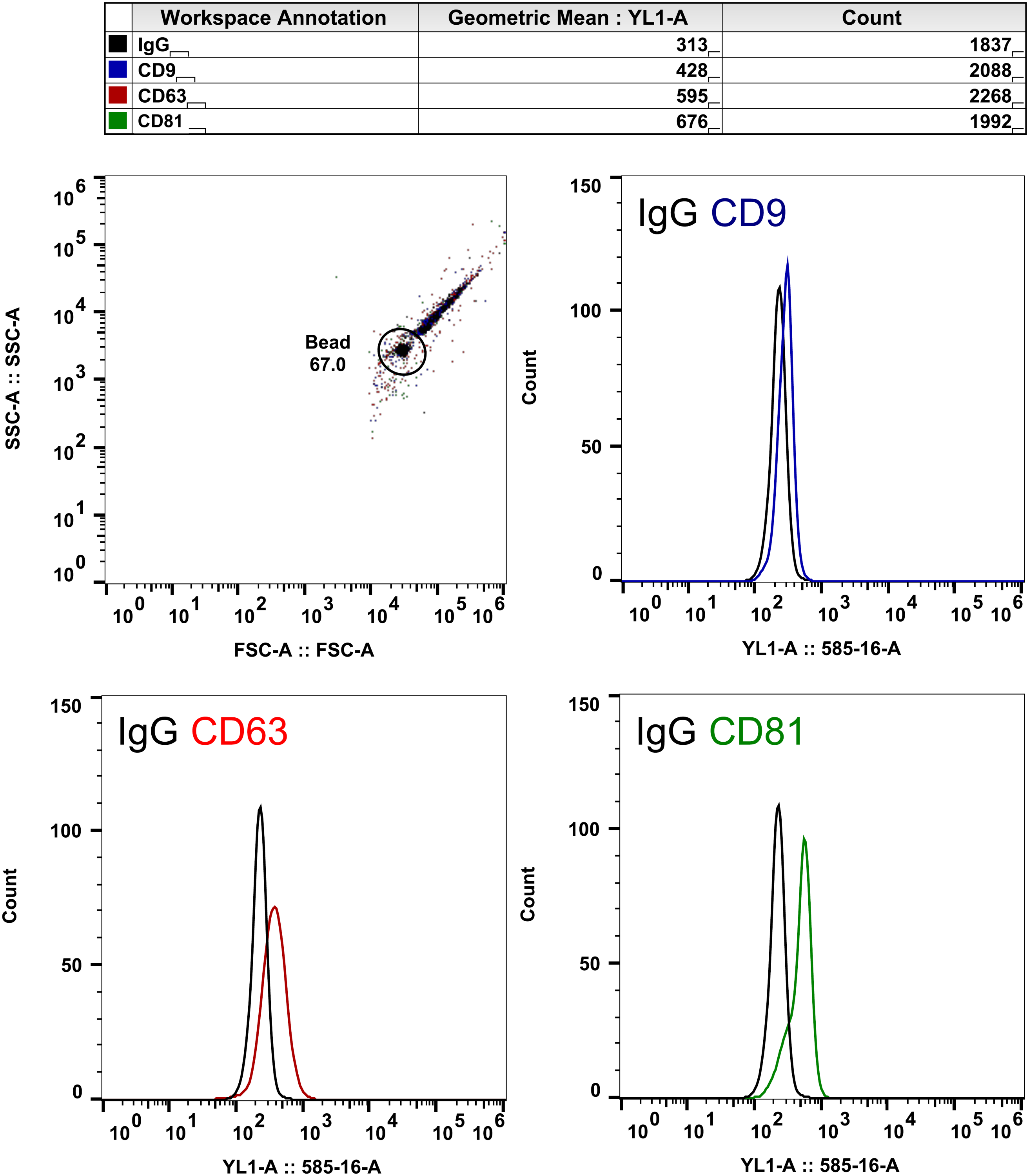

**Supplementary Figure 11. Flow cytometry analysis of A549 extracellular vesicle (EV) tetraspanin markers.** To enable flow cytometry analysis of A549 EVs tetraspanin surface markers, the A549 EVs were bound onto antibody-conjugated beads. Briefly, 100 μl binding samples were setup including 25 μg of A549 cell line EVs (25 μl; #HBM-A549-100; HansaBioMed), 20 μl CD63-antibody conjugated Dynabeads (#10606D, ThermoFisher, USA) and 53 μl PBS (1X).These samples were incubated for 1 h, at 25^o^C with shaking 1000 rpm (Eppendorf, Thermo Mixer C). Samples were washed on a magnetic rack with PBS (1X) and then incubated with 2 μl of either control IgG-PE (#130-113-200), CD9-PE (#130-103-955), CD63-PE (#130-100-153) or CD81-PE (#130-118-342) antibodies (MACS Miltenyi biotec, Germany). Samples were washed on a magnetic rack with PBS (1X), re-suspended in 500 μl PBS (1X) and then analysed using flow cytometry (Attune NxT; YL1-A, Ex 561 nm Em. 585-16 nm). **(Top left)** Forward scatter (FSC-A) and side scatter (SSC-A) contour plot, including the flow cytometry gating strategy (Bead) used for Dynabead analyses. Fluorescence histograms of **(Top right)** CD63+/CD9+, **(Bottom left)** CD63+/CD63+, or **(Bottom right)** CD63+/CD81+ A549 EVs. 1300 events (Dynabeads) were analysed for each sample.

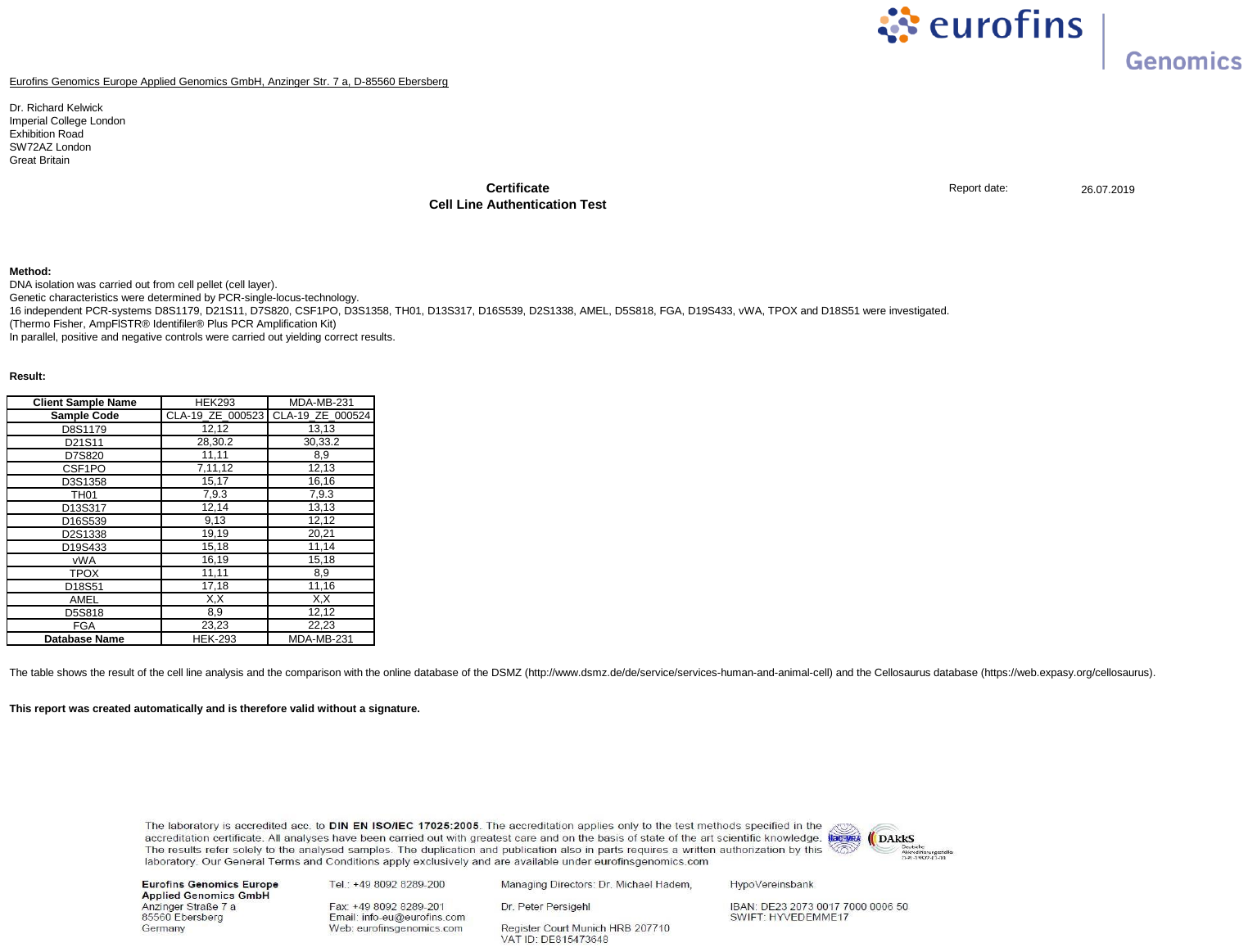

**Supplementary Figure 12. HEK293 cell line validation was carried out by Eurofins scientific (Luxembourg).**
